# The ontogeny of asymmetry in echolocating whales

**DOI:** 10.1101/2022.06.24.497462

**Authors:** Agnese Lanzetti, Ellen J. Coombs, Roberto Portela Miguez, Vincent Fernandez, Anjali Goswami

## Abstract

Extreme asymmetry of the skull is one of the most distinctive traits that characterizes toothed whales (Odontoceti, Cetacea). The origin and function of cranial asymmetry are connected to the evolution of echolocation, the ability to use high frequency sounds to navigate the surrounding environment. Although this novel phenotype must arise through changes in cranial development, the ontogeny of cetacean asymmetry has never been investigated. Here we use three-dimensional geometric morphometric to quantify the changes in degree of asymmetry and skull shape during prenatal and postnatal ontogeny for five genera spanning odontocete diversity (oceanic dolphins, porpoises, and beluga). Asymmetry in early ontogeny starts low and tracks phylogenetic relatedness of taxa. Distantly-related taxa that share aspects of their ecology overwrite these initial differences via heterochronic shifts, ultimately converging on comparable high levels of skull asymmetry. Porpoises maintain low levels of asymmetry into maturity and present a decelerated rate of growth, likely retained from the ancestral condition. Ancestral state reconstruction of allometric trajectories demonstrates that both paedomorphism and peramorphism contribute to cranial shape diversity across odontocetes. This study provides a striking example of how divergent developmental pathways can produce convergent ecological adaptations, even for some of the most unusual phenotypes exhibited among vertebrates.

## 1. Introduction

Skull asymmetry is a rare trait in tetrapods and is usually limited to specific areas of the skull related to sensory abilities, as in some owls displaying asymmetry between the ear canals to allow for directional hearing [1]. Toothed whales (Odontoceti, Cetacea) are one such group with marked directional cranial asymmetry [2], a trait which was acquired in parallel with their ability to echolocate [3]: using high-frequency sounds over 100 kHz to better navigate the complex water environment, locate prey and communicate with co-specifics efficiently [4]. Among mammals, only some bats [5] and a few species of shrews [6] have also convergently evolved the ability to echolocate. This strong directional asymmetry heavily influenced skull evolution in odontocetes [2, 3, 7], including driving unique patterns of phenotypic integration among cranial bones [8].

The ancestors of modern whales, archaeocetes, may have possessed some level of cranial asymmetry in the rostral region to support directional hearing [2, 9]. The other living cetacean clade, baleen whales (Mysticeti), have symmetrical crania, similar to that of the closely related modern ungulates [10], and employ only low frequency sounds for communication [11]. In adult modern odontocetes, asymmetry is mostly prominent in the neurocranium and nasal openings [2, 12]. The bones of the right side of the skull are typically larger and expand leftward, reflecting the morphology of the overlying soft tissues [2, 13], in particular the melon, the fat body that is involved in focusing and transmitting high frequency sounds to the surrounding water, and the phonic lips, valves located in the nasals passages that control sound emission [11, 14]. It has been shown that varying levels of asymmetry in odontocetes are directly correlated with production of sound and its frequency [13, 15], as well as in directionality of hearing [9].

While evolutionary changes in skull asymmetry and its possible drivers have been studied extensively in Odontoceti [e.g. 2, 12, 13], the developmental origin of this trait remains largely unexplored, despite the established importance of ontogenetic shifts in the rise of other unique cetacean traits, such as hind limb reduction or tooth loss [16, 17].

Here, we address this gap with a quantitative analysis to answer the question: how does skull asymmetry develop in toothed whales?

We approach this question by applying landmark-based 3D geometric morphometrics (GM) methods in a dataset that represents all prenatal and postnatal growth stages of an ecologically and taxonomically diverse selection of modern odontocetes: Delphinidae (oceanic dolphins – *Stenella* and *Lagenorhynchus*, pilot whales – *Globicephala*), Monodontidae (beluga – *Delphinapterus*), and Phocoenidae (porpoises – *Phocoena*). With these data, we characterize both degree and pattern of asymmetry and skull shape through ontogeny to understand the role of ecological adaptations and phylogenetic relationships in determining the development and evolution of skull asymmetry.

## 2. Results

### (a) The ontogeny of asymmetry

Cranial asymmetry was quantified through ontogeny as the overall mean Euclidean distance between directional asymmetric (DA) and symmetric shape, after excluding the variably present interparietal bone [18] (Materials and Methods). Asymmetry starts at relatively low levels in the early fetal stages of all taxa and is closely aligned with phylogenetic relationship, with *Delphinapterus* and *Phocoena* sharing a more symmetric cranial shape relative to the three species of Delphinidae (figure 1, electronic supplementary material, figure S1, table S1). Ontogenetic trajectories of asymmetry then diverge markedly. Asymmetry increases particularly rapidly between the early fetal stages and birth in *Delphinapterus*, while *Phocoena* maintains significantly lower levels for its entire growth, although in both taxa the only significant increase in asymmetry levels is in the prenatal growth phase and then they stabilize postnatally. This steep increase allows *Delphinapterus* to reach high levels of asymmetry comparable to *Globicephala* in the juvenile stage. The three sampled species of Delphinidae increase asymmetry more gradually, with significant changes only occurring in the postnatal stages in *Lagenorhynchus* and *Globicephala*. Among adult specimens, the three delphinids (*Lagenorhynchus, Stenella, Globicephala*) and the monodontids (*Delphinapterus*) displayed significantly higher levels of asymmetry than the phocenid (*Phocoena*). *Globicephala* also presents a more asymmetric skull than the other oceanic dolphins (*Lagenorhynchus*, *Stenella*). These asymmetry levels found in the adults are consistent with previous studies [2, 13], and validate the methodology used here to quantify cranial asymmetry.

**Figure 1.**
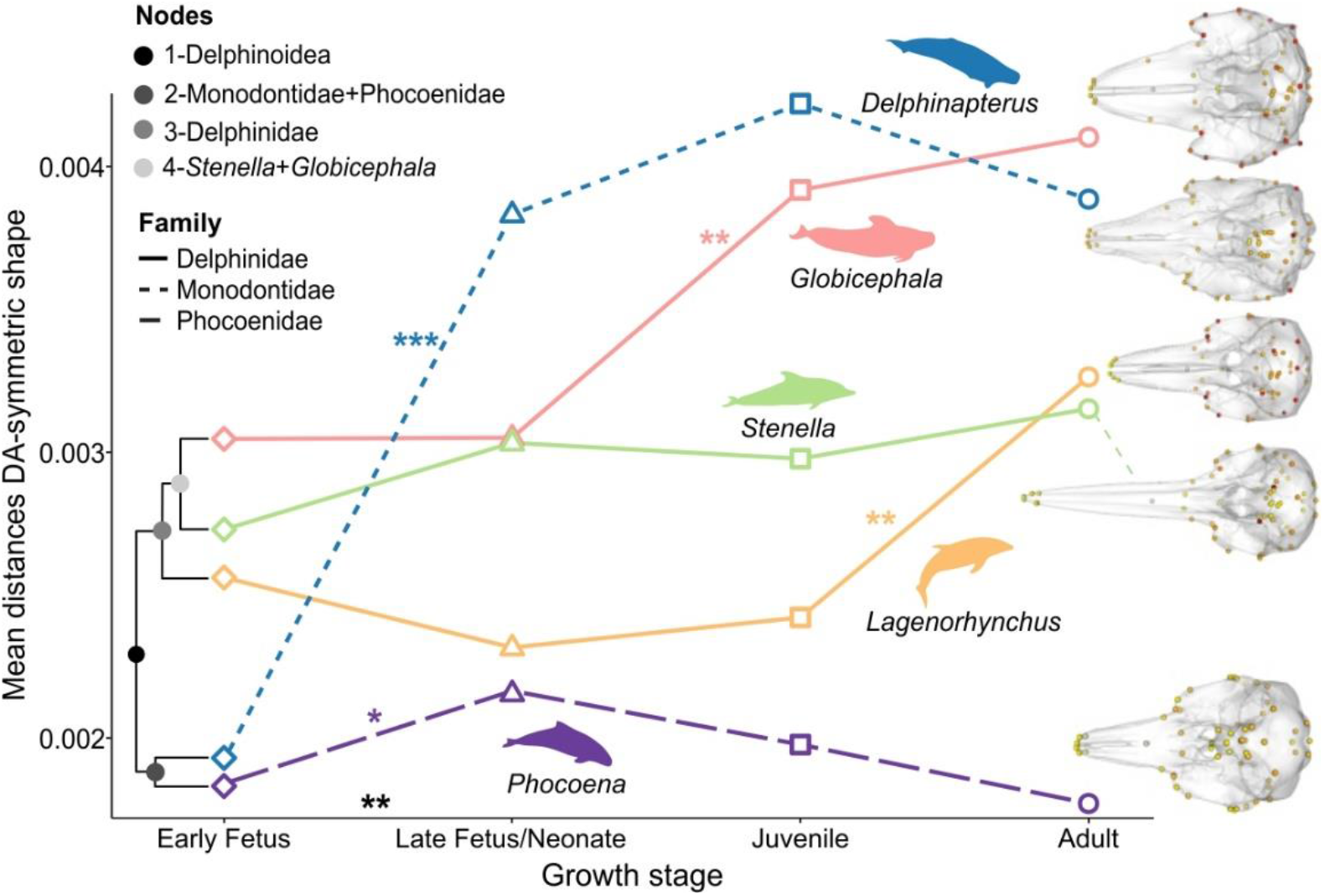
Ontogeny of cranial asymmetry in toothed whales. Degree of asymmetry is quantified as the mean distance between the directional asymmetric (DA) and symmetric shape across all landmarks for all specimens of the same genus at each growth stage. The phylogeny drawn on the left highlights the close correspondence between phylogenetic structure and starting level of asymmetry in ontogeny. The skulls on the right represent variation in asymmetry in adult skulls. Landmarks in darker colours indicate more asymmetric regions, as they have a larger distance between the DA and symmetric shape relative to mean of all adults. Significant shifts in asymmetry levels between growth stages in the entire dataset and per genus are indicated by asterisks (‘***’ p<0.001, ‘**’p<0.01, ‘*’ p<0.05). See electronic supplementary material, table S1, figure S1-S2 for full results.

The distribution of asymmetry across the skull is highly divergent among taxa from the late fetal stages (electronic supplementary material, figure S2). The low variation in *Phocoena* is distributed throughout the cranium, and in closely related *Delphinapterus* is mostly concentrated in the supraoccipital and nasal region, with a visible leftward shift of the skull midline (figure 1). Delphinidae instead have high levels of variation in the neurocranium, from the antorbital notches to the supraoccipital, with *Stenella* displaying a prominent asymmetry in the dorsal end of the premaxilla. Therefore, while *Delphinapterus* and *Globicephala* present similar asymmetry levels in the adults, the different distribution of the asymmetry still reflects their different evolutionary history, as does their markedly different levels of asymmetry in early ontogeny.

### (b) The ontogeny of skull shape

Phenotypic trajectory analysis [PTA, 19] of skull shape ontogeny reinforces the morphological convergence between *Delphinapterus* and *Globicephala* and also highlights the peculiarity of skull development in *Phocoena* (figure 2, electronic supplementary material, figure S3, table S2). The phenotypic trajectories of all taxa are similar in shape but have unique directions. *Delphinapterus* and *Globicephala*, the two most asymmetric genera, are characterized by a longer trajectory compared to the rest of the dataset. However, the distribution of shape change in these two taxa is very different, owing to their phylogenetic history. *Delphinapterus* has a very prominent shape change early in the ontogeny followed by shorter shifts among postnatal stages, while *Globicephala* has similar amounts of shape change across all growth stages. The other delphinids have distinctly different trajectories that do not overlap with *Globicephala*, though they also display consistent shifts in shape across developmental stages. *Phocoena* has a unique phenotypic trajectory, mostly following a straight line along PC1 and with a very small shape change between juvenile and adult stages. This is likely connected to its low and constant levels of skull asymmetry.

**Figure 2.**
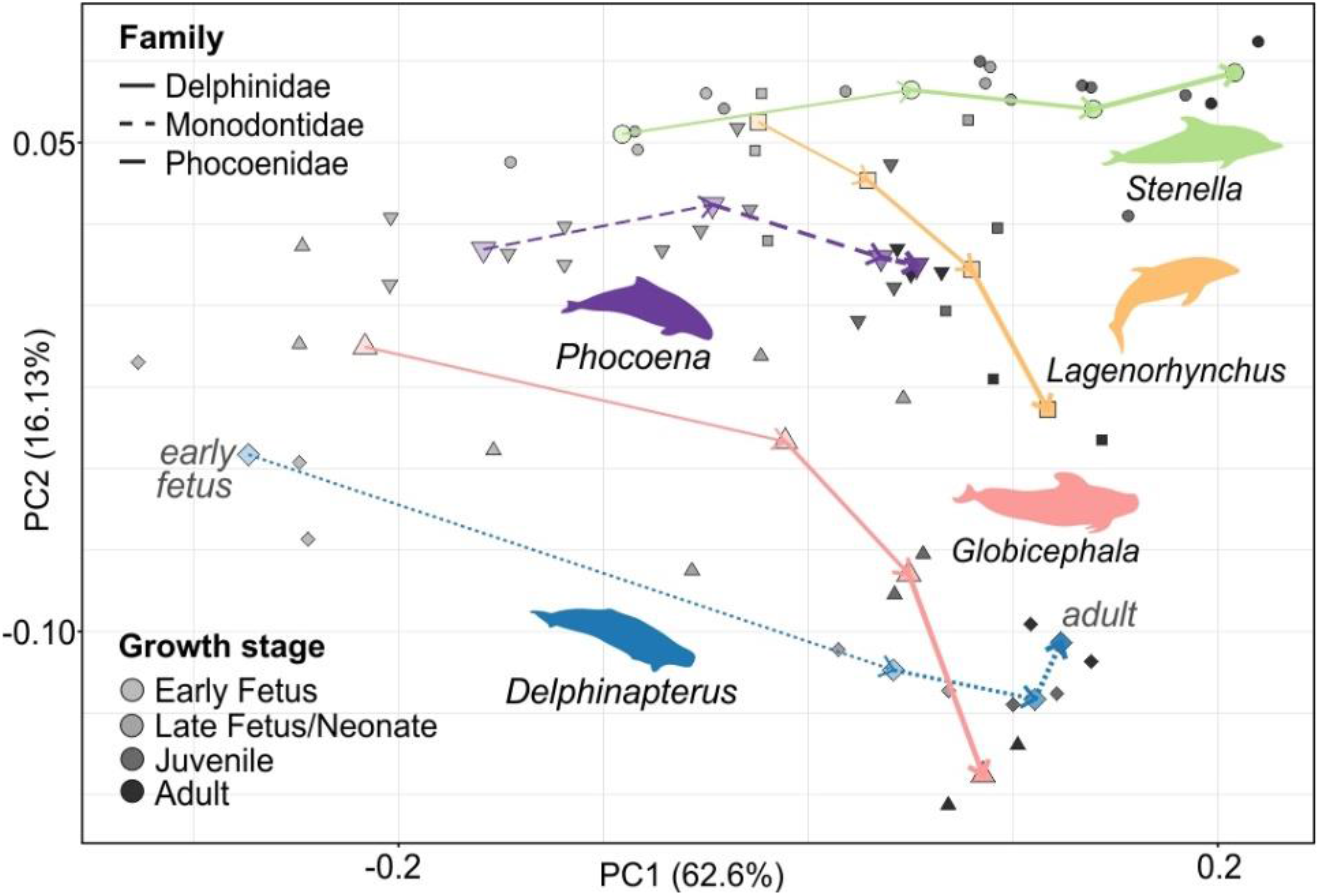
Trajectory of skull shape growth (PTA) in each taxon. Direction of ontogeny follows PC1 axis, from negative (early fetal stages) to positive values (adults). *Delphinapterus* and *Globicephala* display longer trajectories than the other taxa. See electronic supplementary material, table S2, figure S3 for full results.

### (c) Evolution of skull allometry

Another important aspect of ontogeny is allometry, or the rate of shape change relative to size [20]. When studying ontogenetic allometry, it is possible to not only characterize the trends observed in modern taxa, but also to estimate the allometric trajectories in the ancestors [21] and in turn determine the direction of heterochronic change between the ancestral nodes and the descendants [20]. This allows direct testing of whether the patterns observed in ontogeny are due to the influence of phylogeny or are instead brought about by ecological convergence. Since we found differences in the prenatal and postnatal rate of change in both shape and asymmetry, we first tested if allometric trends were also significantly different in these two phases of ontogeny, as has been proposed in other mammals [22] and amniotes [23]. Our analysis identified a model with one varying break point and with different slopes for each taxon as the preferred model (electronic supplementary material, table S3). Within each genus, we found significant absolute distances in allometric slope and length differences between prenatal and postnatal stages in all three Delphinidae (electronic supplementary material, figure S4*a*, table S4). Allometric growth in Monodontidae and Phocoenidae does not show a significant change before and after birth.

In comparing allometric slopes among genera, *Delphinapterus* and *Globicephala* are significantly different to each other in all metrics during prenatal growth, but not postnatally (electronic supplementary material, figure S4*b*, table S4). Additionally, both taxa differ significantly from *Phocoena* and *Stenella* in prenatal ontogeny, while only *Globicephala* retains a significant difference with *Stenella* postnatally. This result is consistent with previous observations that the level of convergence between *Delphinapterus* and *Globicephala* increases progressively during growth. *Phocoena*, while exhibiting low levels of asymmetry and shorter postnatal growth, does not exhibit a unique allometric trend compared to the other taxa.

To assess which heterochronic processes generated these differences in allometry among the sampled taxa, we estimated the ancestral slopes and intercepts for prenatal and postnatal growth in the following ancestral nodes: 1) the common ancestor of Delphinoidea, the group that includes all the families analysed, 2) the ancestor of Phocoenidae and Monodontidae, 3) the ancestor of Delphinidae, and 4) the common ancestor of *Globicephala* and *Stenella*, within Delphinidae. The regression plot including both modern and ancestral allometric regressions shows a strong separation between *Globicephala* and *Delphinapterus* and all other taxa, both prenatally and postnatally (figure 3*a*, electronic supplementary material, figure S5*a*-*b*). These taxa have consistently lower rates of development. *Phocoena* instead presents a relatively conserved allometric trend prenatally, which closely tracks its ancestral node, while postnatally it is has a lower rate.

**Figure 3.**
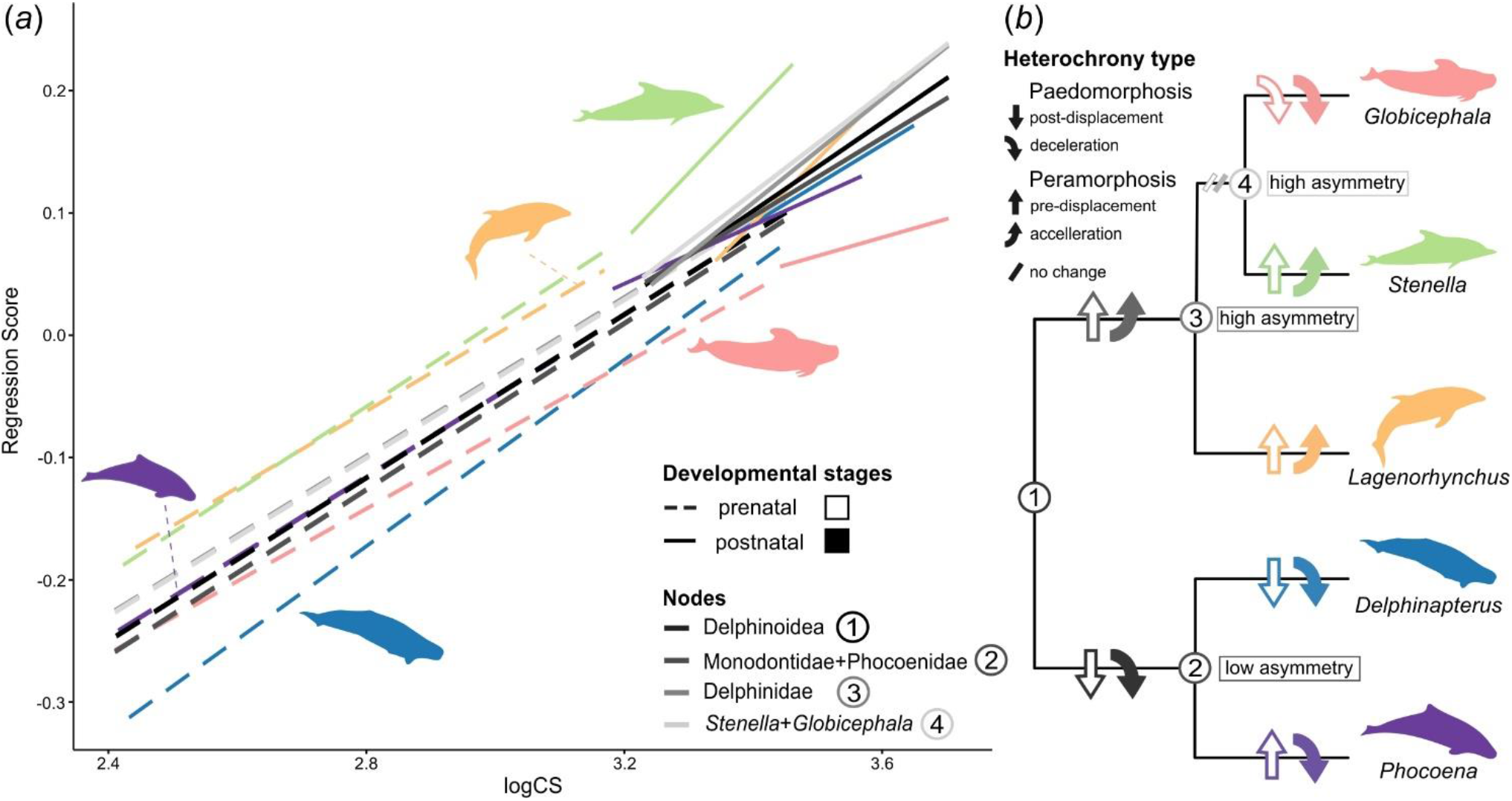
Allometric patterns in prenatal and postnatal ontogeny of toothed whales’ taxa and their direct ancestors. (*a*) Regressions of skull shape and size (logCS); (*b*) Phylogeny with inferred direction of heterochronic change for each node. Similar lower slopes in *Delphinapterus* and *Globicephala* were acquired independently, due to a major paedomorphic shift in *Globicephala* from its peramorphic ancestor. *Phocoena* prenatal slope overlaps the ancestor of Delphinoidea (node 1), while postnatally a paedomorphic shift has occurred. Degree of asymmetry at each node is also reported on the phylogeny. Regressions are represented separately for the prenatal and postnatal stages in electronic supplementary material, figure S5*a*-*b* to better visualize difference between taxa and reconstructed ancestral nodes. For significance patterns see electronic supplementary material, figures S4*a*-*b*-S6*a*-*b*.

The only significant difference recovered between prenatal and postnatal allometry in ancestral slopes is in node 2 (ancestor of Phocoenidae and Monodontidae). Evolutionary changes in allometry are concentrated in prenatal growth, with most ancestral prenatal slopes are different from each other and from modern taxa, with the notable exception of *Phocoena*. In postnatal ontogeny, in contrast, only *Stenella* differs significantly from the ancestral nodes (electronic supplementary material, figure S6*a*-*b*, table S5).

Combined, these observations allow us to characterize the direction of heterochronic changes that have occurred during the evolution of these taxa (figure 3*b*). Paedomorphism, which is defined as the retention of juvenile ancestral traits in the adult of the descendant [20], is recognizable by a decrease in slope (deceleration) or intercept (post-displacement) in an extant taxon relative to its ancestor. This type of heterochrony is recognizable in the evolution of Phocoenidae and Monodontidae, in both prenatal and postnatal ontogeny. This process is particularly visible in *Delphinapterus*, while *Phocoena* only presents deceleration in postnatal growth. Convergently, *Globicephala* also shows decelerated growth relative to its ancestor both prenatally and postnatally. In fact, the opposite heterochronic transformation, peramorphism, defined as the presence of adult ancestral traits in the juvenile of the descendant [20], is present in Delphinidae. An increase in intercept value (pre-displacement) is visible prenatally in the ancestor of Delphinidae and in the other two oceanic dolphins, *Stenella* and *Lagenorhychus*. Postnatal ontogeny of these same taxa and nodes is instead characterized by higher slope values (acceleration). This estimate suggests that the evolution of convergent skull morphologies in *Delphinapterus* and *Globicephala* occurred independently via parallel changes in different aspects of development: an increase in asymmetry levels in the first taxon and a paedomorphic shift in shape development in the second. These changes were likely driven by similar ecological pressures. The unique phenotypic trajectory and ontogeny of asymmetry of *Phocoena* instead are likely derived from its ancestral condition.

## 3. Discussion

### (a) Shared ecology drives convergence in skull ontogeny

Closely related species display similar levels of asymmetry early in ontogeny, but asymmetry in adults shows clear association with ecology, regardless of phylogeny. This convergence is achieved through divergent ontogenetic trajectories in cranial asymmetry, which is most evident in comparing *Delphinapterus* with the closely related *Phocoena* and the distantly related, but ecologically similar, *Globicephala*. Both *Delphinapterus* and *Phocoena* start off with low levels of asymmetry but, while *Phocoena* remains at a similar level throughout ontogeny, *Delphinapterus* has a steep increase in asymmetry prenatally to reach similar levels to *Globicephala* (figure 1). *Delphinapterus* and *Globicephala* also share other aspects of their skull ontogeny: they present similar phenotypic trajectories (figure 2) and convergently acquired paedomorphic allometric growth (figure 3).

The clades including *Delphinapterus* and *Globicephala* diverged approximately 20 million years ago (Mya) [24] and have since convergently evolved similar ecological adaptations. They occupy non-overlapping environments and geographical ranges, with *Globicephala* found worldwide in tropical and temperate waters [25], while *Delphinapterus* only lives around the ice pack in the North Pole [26]. Nonetheless, they share remarkable similarities in feeding strategy and prey selection. Both taxa are primarily suction feeders, with a large part of their diet coming from squid and small fish, with the addition of benthic invertebrates for the *Delphinapterus* [25–27]. This is reflected in their shared cranial morphology, with a broader and shorter rostrum compared to other taxa [28, 29]. To locate and capture their prey, they both rely heavily on echolocation. These taxa employ similar broadband (BB) hearing frequencies, spanning from ~10 to 120 kHz, with peak sensitivity at mid-range between 40 and 80 kHz [25, 26]. This results in comparable levels of skull asymmetry observed in the adults [2] and during the postnatal stages of development, as well as in overlapping inner ear morphologies [30]. They also share similar body size of about 5 m in the adults [25, 26], which has been found to correlate with echolocation frequency [31] and prey size [29]. It has been hypothesized that the hearing frequency used by *Delphinapterus* and the other extant genus of Monodontidae (*Monodon*, narwhal) are a key adaptation for navigating their complex polar environment, avoiding ice and finding prey in murky waters [30, 32, 33]. *Globicephala*, unlike other members of Delphinidae, tends to forage at night during deep dives of over 100 meters below the surface, a strategy shared with other toothed whales such as Physeteriidae (sperm whales) [25, 34]. It is possible that the same frequencies used by Monodontidae help *Globicephala* avoid obstacles and find prey in the dark. Another aspect of ecological convergence is the sociality of these two taxa. They both live in large groups spanning from 20 to a few hundred individuals, mostly composed of females and related juveniles, with males living separately and only occasionally joining the larges pods [25, 26]. Due to this behaviour, they retain whistles and other tonal sounds, acoustic features commonly found in species of toothed whales that form complex social groups [35].

This impressive convergence in ecology and connected morphological traits in *Delphinapterus* and *Globicephala* is achieved through parallel but different ontogenetic shifts, which are able to overwrite phylogenetic patterns, as has been shown in birds [36] and crocodiles [21, 23]. A major peramorphic shift in skull asymmetry and shape development in prenatal ontogeny of *Delphinapterus* allowed this taxon to evolve its relatively large and asymmetric skull, while an overall deceleration in rate of skull growth in *Globicephala* likely played an important role in the evolution of its distinct morphology among Delphinidae. The anatomical requirements connected to the use of specific echolocating frequencies are likely the main driver behind this phenomenon. Their distant ancestry is still recognizable in the levels of asymmetry in the early fetal stages and in its pattern of change through ontogeny, as well as in the phenotypic trajectory of *Delphinapterus*.

### (b) Retention of skull symmetry in ontogeny of Phocoenidae

Among Odontoceti, Phocoenidae are commonly described as having immature traits, with a proportionally shorter rostrum and larger braincase compared to delphinids with a similar feeding mode, such as *Lagenorhynchus* [37, 38], and lower levels of asymmetry [2, 13]. *Phocoena* is the only taxon in the dataset to maintain low levels of asymmetry throughout its ontogeny, with early fetuses and adults sharing mostly symmetric skulls (figure 1), a characteristic also reflected in their soft tissue morphology [37]. The closely related *Delphinapterus* starts from similarly low levels of asymmetry but goes on to develop a highly asymmetric skull in adults. Despite drastically different levels of asymmetry and phylogenetic distance, *Phocoena* displays a similar cranial morphology to oceanic dolphins like *Stenella* and *Lagenorhynchus* throughout development (electronic supplementary material, supplemental results, figure S7*a*-*b*, table S6). *Phocoena* and *Delphinapterus* also share a similar shortened postnatal development of skull shape (figure 2), as well as a generally paedomorphic trend in allometry inherited from their ancestor (figure 3). Overlapping feeding modes and prey type might be driving the apparent morphological similarities in overall skull shape and growth trajectory between *Phocoena* and some longirostrine oceanic dolphins, but these shared ecological traits do not explain the retention of low levels of skull asymmetry in adults and of other unique characteristics of their development. The unique echolocating abilities of Phocoenidae are the most likely limiting factor in the evolution of this group, and they influence multiple aspects of their ontogeny.

The majority of the 75 extant species of odontocetes utilize broadband (BB) clicks in the 20-150 kHz for locating prey and communicating, and they present typically high levels of cranial asymmetry, as is the case in the oceanic dolphins present in this dataset and *Delphinapterus*. But Phocoenidae and three other distinct groups, the pygmy and dwarf sperm whales (*Kogiia* spp.), the franciscana river dolphin (*Pontoporia blainvillei*), the Lissodelphininae (*Cephalorhynchus* spp., *Lissodelphis* spp.), have convergently evolved the use of narrow-band high-frequency sounds (NBHF) [4]. These taxa have higher sensitivity at the upper end of the hearing spectrum (125-140 kHz) and selectively use these frequencies to communicate rather than lower ones like BB species [4, 13]. It has been hypothesized that ecological pressures have driven the convergent evolution of NBHF. They prevalently inhabit coastal and riverine environments, with the exception of Kogiidae [4]. Using NBHF sounds might be advantageous for navigating the complex environment that surrounds them [39], and it also allows these taxa to avoid predation by large predators such as killer whales, which do not hear sounds at such high frequencies [4, 39]. Nevertheless, they employ a variety of feeding strategies, from the specialized suction feeder *Kogia* to the raptorial feeder *Pontoporia* [4, 39, 40]. Morphologically, while they convergently evolved similar inner ear shapes in order to use this specialized type of echolocation [4], they do not consistently display similar skull traits. NBHF taxa tend to retain a largely symmetric skull, similar to that observed in early developmental stages of modern odontocetes [37], as well as in stem taxa [3]. However, this is far from a consistent pattern, with some taxa presenting high levels of asymmetry both in the neurocranium and correlated soft tissues [13, 41]. They also have different rostral morphologies, as this characteristic is tightly linked with feeding strategy in odontocetes [7, 28, 29]. The only trait that they all share is a relatively small body and skull size [4], which has been directly linked to the ability to hearing these high frequency sounds due to scaling of the melon [31].

As *Phocoena* is the only NBHF taxon in the dataset, its unique ontogeny might be associated with the evolution of this specialized hearing frequency. A shortened and slower postnatal shape growth are the key developmental features that allow adult Phocoenidae to use NBHF sounds, allowing for the correct scaling of the melon relative to its body size [31]. Retention of skull symmetry is likely a product of these other aspects of ontogeny, rather than necessary to develop the ability to use NBHF. This would also explain the asymmetric skull observed in some species of Lissodelphininae [42] and Kogiidae, one of the groups of Odontoceti with the highest skull asymmetry [2, 13], as well as the asymmetric soft tissue morphology of *Pontoporia* [37, 41]. Immature features of Phocoenidae have been attributed to paedomorphic shifts in development occurring in the evolution of this group [37, 38]. Based on the reconstructed allometric trajectories and the development of skull asymmetry in closely related *Delphinapterus*, it is likely that they have instead inherited both skull symmetry and slower and shorter development from the ancestor of the group, rather than secondarily acquiring them through paedomorphic changes in connection to NBHF specifically. However, heterochrony might play a role in the evolution of NBHF hearing in other taxa such as Kogiidae and their ontogeny should be studied in detail to test this hypothesis.

### (c) Multiple ontogenetic routes to asymmetry

Given the clear divergences observed in the ontogeny of skull shape and asymmetry observed here, data from fossil taxa may elucidate when and why these shifts occurred in the evolution of toothed whales. Cranial asymmetry appears in the earliest toothed whales Xenorophidae† in the mid Oligocene (~30 Mya), likely in connection with the evolution of echolocation [2, 3]. Xenorophids† presented a variety of ecological adaptations, from raptorial feeding to specialized suction feeding, reflected in their disparate skull morphologies with varying degrees of telescoping of the neurocranium [43]. As asymmetry levels and degree of telescoping tend to increase gradually during ontogeny in the modern taxa we examined, we can hypothesize that these stem odontocetes had a generally slower or shorter growth compared to extant odontocetes. A medium degree of cranial asymmetry is retained in most odontocetes after this initial shift, with notable increases in Platanistidae, which include the modern South Asian river dolphin (*Platanista gangentica*), and in extant and extinct sperm whales (Physeteriidae and Kogiidae) [2, 43]. These occurrences of higher asymmetry may signal the evolution of convergent ecological adaptations [2, 4, 13, 41] though achieved through different developmental routes, as it is the case in *Delphinapterus* and *Globicephala* in the present dataset. Qualitative studies of the ontogeny of Physeteriidae suggest that they present high levels of asymmetry from the early ontogenetic stages [44], providing additional support for the hypothesis that phylogeny influences the starting level and patterning of asymmetry in ontogeny. If these similar levels of asymmetry were also present in the early fetal stages of closely related Kogiidae, this would explain why they present a highly asymmetrical skull while using NBHF sounds. On the other side of the spectrum, the fossil hyper-longirostrine group Eurhinodelphinidae† presents an overall symmetric skull [2, 45]. This apparent reduction of skull asymmetry might have been coupled with the retention of asymmetrical nasal soft tissues and melon as observed in Pontoporiidae, as this taxon also presents an elongated rostrum [41].

Another shift occurs at the base of Delphinoidea (Phocoenidae + Monodontidae + Delphinidae) in Kentriodontidae†, a likely paraphyletic fossil group of toothed whales from the early Miocene (~20 Mya) [2, 46]. These taxa have overall symmetrical crania, and the type species *Kentriodon pernix†* was reconstructed as having used NBHF sounds based on the morphology of its inner ear [2, 4, 37]. It is possible that Phocoenidae and Monodontidae retained their low levels of asymmetry in the early fetal stages from a common ancestor closely related to this extinct group. In fact, *Odobenocetops†*, a fossil taxon with limited echolocating abilities and a mostly symmetric skull, is hypothesized to be a close relative of modern Monodontidae [2]. Therefore, the most parsimonious hypothesis, supported also by our results, is that Monodontidae independently acquired highly asymmetric skulls adapted to BB frequency hearing from an ancestor with a relatively symmetrical cranium and that employed NBHF sounds [2]. This was possible via a shift in rate of asymmetry and overall skull ontogeny in the prenatal stages. *Phocoena* instead retained the ancestral ecology and NBHF hearing frequencies, with the associated pattern of asymmetry and cranial development.

Fossil and early diverging Delphinidae share moderately high levels of skull asymmetry, most prominent at the dorsal end of the premaxilla, as exemplified by *Eodelphis†* and *Orcinus* (killer whale) [2, 47], similar to what is observed in *Stenella*. Both *Lagenorhynchus* and *Stenella* retain a level of asymmetry likely comparable to their ancestor, which only increases slightly during ontogeny. A shift towards higher asymmetry occurred in the ancestor of the subfamily Globicephalinae, to which *Globicephala* belongs, along with other highly asymmetric taxa such as *Pseudorca* (false killer whale) [2, 7]. It is possible that other genera in this subfamily also have comparatively paedomorphic allometric growth relative to other Delphinidae, while achieving higher levels of skull asymmetry via a longer phenotypic trajectory.

## 4. Conclusions

This first quantitative analysis of cranial shape development in toothed whales demonstrates that a strong phylogenetic signal in early ontogeny is overwritten by ecologically-driven transformations occurring at different stages of growth. A peramorphic shift in prenatal ontogeny of asymmetry of *Delphinapterus* has allowed this taxon to obtain highly asymmetric skulls, similar to that of the delphinid *Globicephala*, while maintaining a paedomorphic growth trend for skull shape. *Globicephala* instead has secondarily slowed its cranial development while maintaining high levels of asymmetry. Ecological factors such as feeding and sociality in the two distantly related taxa have probably driven this convergence, shaping their echolocating abilities and body size [30, 31]. *Phocoena* has uniquely low skull asymmetry and truncated postnatal ontogeny compared to the other sampled species. Based on observations of modern and fossil taxa, this pattern is likely retained from the ancestral state of the clade, with additional paedomorphism in postnatal ontogeny being responsible for the juvenile traits and small body size observed in the adults [48]. This paedomorphic tendency is likely correlated with the evolution of NBHF hearing through scaling of the skull and melon in Phocoenidae and other taxa that use these frequencies for echolocation [31]. Retention of skull symmetry is a by-product of this lower developmental rate and reflect the ancestral state of Phocoenidae, as it is not consistently present across taxa that employ NBHF sounds [4, 37]. *Stenella* and *Lagenorhynchus* instead show convergent peramorphic patterns of allometry and similar level of skull asymmetry, possibly inherited from the ancestor of Delphinidae [2].

Even in a dataset composed of relatively closely related taxa, developmental patterns display large variations. These changes can increment or negate each other through complex interactions in different aspects of development [20]. Studying them is key to better understanding of adaptive patterns of evolution and distinguishing between traits that are directly related with functions and behaviours and those that are a remnant of the phylogenetic history of the clade [16]. Investigating ontogenetic changes in Cetacea has already contributed to reformulate hypotheses on the evolutionary origin of some of their most distinctive traits such as hind limb loss [e.g. 16, 49], teeth simplification and transition to baleen [e.g. 17, 50], ear bone morphology [51], and, here, skull asymmetry. The role of ontogeny in the evolution of unique morphological traits and ecological adaptations remains largely unexplored in many lineages, and we hope that similar approaches to one used here can be applied to future studies in the revived field of evo-devo.

## 5. Materials and Methods

### (a) Morphometric data

The dataset is composed of 58 specimens representing all phases of growth of five odontocete genera from three families: *Delphinapterus* (beluga – Monodontidae), *Phocoena* (common porpoises – Phocoenidae), *Globicephala* (pilot whales – Delphinidae), *Lagenorhynchus* (white-beaked dolphin, white-sided dolphins – Delphinidae), *Stenella* (spinner dolphins, spotted dolphins – Delphinidae). While some of these genera are comprised of multiple species, their morphological similarity and the large phylogenetic distance between genera allows us to use these genera as taxonomic units in the analysis [24] (electronic supplementary material, supplemental methods, dataset S1).

Skull morphology was digitized using computed tomography (CT) scanning for fluid preserved specimens and disarticulated osteological specimens (electronic supplementary material, supplemental methods, dataset S2) and using a hand-held surface scanner (Creaform GoSCAN!) for whole osteological specimens. 3D rendering of CT data was performed in Avizo Lite. All meshes were imported in Geomagic Wrap 2017 for cleaning, reduced to 1.5 million faces, and then exported as PLY for landmarking and visualization (electronic supplementary material, supplemental methods).

To quantify the shape of the skull, 64 single points landmarks and 43 curves were placed on the surfaces using Stratovan Checkpoint. All points and curves were digitized manually for the right and left side of the cranium to capture asymmetry (electronic supplementary material, supplemental methods, figure S8, table S7). The coordinates for each specimen were exported in PTS format to be analysed.

### (b) Data analysis

The shape coordinates for all specimens were imported in R [52] for data analysis. After resampling and sliding the curve semilandmark and fixing missing and absent points (electronic supplementary material, supplemental methods), the final dataset used for analysis is composed of 462 points, of which 64 are fixed landmarks and 398 are semilandmarks. Procrustes superimposition (GPA) was performed using the ‘gpagen’ function in ‘geomorph’ v. 4.0.1 [53] to align and scale all specimens and extract Centroid Size (CS) to be used a measure of skull size in allometry analysis. Taxonomic and age information were separately imported to use as covariates. The specimens were binned into four growth categories based on their approximate age and total length: early fetus (from conception to <50% gestation), late fetus and neonate (from >50% gestation to birth, up to the first year of life), juvenile (after the first year until sexual maturity), adult (after sexual maturity) (electronic supplementary material, supplemental methods). Creating these broad age bins allowed to account for possible errors in age and length estimation and created an equal distribution of the dataset between taxa and growth categories, with at least two specimens per stage for each taxon (electronic supplementary material, dataset S1).

#### (i) Asymmetry analysis

Asymmetry levels were assed only using fixed points landmarks, as using semilandmarks is not implemented consistently in the available methods [2]. We used the function ‘bilat.symmetry’ from the ‘geomorph’ package to extract the Directional Asymmetric (DA) shape component and symmetric component of skull shape for each specimen. We then calculated the Euclidean distance between each of the symmetric shape landmarks and the DA shape landmarks was calculated with the ‘linear.dist’ function in the package ‘landVR’ v. 0.4 [54]. We used these data to calculate mean distances for each landmark averaging specimens of the same taxon binned in the same growth stage, revealing information on where asymmetry is most visible in the skull. To compare the overall level of asymmetry in the skull across categories and taxa and identify significant differences, we used the ‘t.test’ and ‘pairwise.t.test’ functions from the ‘stats’ package v. 4.1.1 [52]. This also produced the mean distance for all the landmarks for each stage and taxon, which is useful to visualize how asymmetry varies in the dataset. As the interparietal bone is variably present in adult odontocetes [18, 55], we decided to remove the three landmarks marked on this bone from the asymmetry analysis presented in the main text (figure 1). However, we did perform the same full set of analyses on the whole skull configuration, and they produced comparable results (electronic supplementary material, supplemental methods, figure S9-S10-S11, table S8).

#### (ii) Shape variation

We assessed the variation of shape in the dataset using the whole configuration including semilandmarks by performing a PCA with the ‘gm.prcomp’ function (electronic supplementary material, supplemental methods) and a phenotypic trajectory analysis across the ontogeny of each taxon with ‘trajectory.analysis’ [19], both implemented in ‘geomorph’. The analysis of trajectory of shape change through ontogeny for each genus was performed on the raw shape coordinates. The function used also allows to significantly test pairwise differences between trajectories.

#### (iii) Allometry analysis

As it has been proposed in other mammals [22], we first tested if the prenatal and postnatal allometries were significantly different from each other using the package ‘mcp’ v. 0.3.0 [56] following [23]. This package calculates the position of a breakpoint in a regression slope and allows us to test if the model with changing slopes is better at explaining the variation than a common slope model. We found that tested models with one break point were consistently preferred (electronic supplementary material, supplemental methods, figure S12*a*-*b*, table S4). Given the data binning chosen for the ontogenetic stages, we proceeded to create and test differences of allometry between taxa in the prenatal (early fetus and late fetus/neonate categories) and postnatal (juvenile and adult categories) ontogeny. We used ‘procD.lm’ in ‘geomorph’ with specimens divided by genus and stage of growth (prenatal and postnatal) to reconstruct the allometric regressions, and we tested pairwise differences between slopes using ‘pairwise’ in the package ‘RRPP’ v. 1.1.2 [57]. We then extracted the regression parameters (slopes and intercepts) and used them to reconstruct ancestral allometric trends, a key step to assess the heterochronic processes at play [20], adapting the methodology presented in [21]. Using a simplified phylogeny with branch lengths extracted from [24], we calculated ancestral slope and intercepts parameters for the nodes using the ‘fastAnc’ function in ‘phytools’ v. 1.0 [58]. To assess significant changes in the allometric slopes between ancestral nodes and extant taxa, we used the package ‘emmeans’ v. 1.7.2 [59] (electronic supplementary material, supplemental methods). Graphics of plots were improved using the ‘ggplot2’ package v. 3.3.5 [60].

## Supporting information

electronic supplementary material

## Data accessibility

The surface files will be uploaded to the open access repository Phenome10K (https://www.phenome10k.org/) after publication. Landmarks, classifiers, and other data necessary for analyses, as well as all the code used, are uploaded to GitHub https://github.com/AgneseLan/ontogeny-asymmetry-dolphin (https://doi.org/10.5281/zenodo.6553285).

## Author Contributions

A.L. and A.G. designed the study, A.L. and E.J.C. developed the methodology, A.L. performed the analysis and wrote the manuscript, R.P.D. and V.F. provided resources and support for data collections and curation, E.J.C., R.P.D., V.F. and A.G. reviewed and edited the manuscripts.

## Acknowledgments

Without natural history collections and curators, this study would have not been possible. We would like to thank Brett Clark from the Natural History Museum, London, UK (NHMUK) for the help with CT scanning specimens from the NHMUK. For sharing CT data of specimens from the Smithsonian Institution in Washington, D.C. (USNM), we thank the Michael McGowen and Janine Hinton.

## References

[1] Norberg, R.A. 2002 Independent evolution of outer ear asymmetry among five owl lineages; morphology, function and selection. In Ecology and conservation of owls (eds. I. Newton, R. Kavanagh, J. Olsen & I. Taylor), pp. 329–342. Clayton, CSIRO Publishing.

[2] Coombs, E.J., Clavel, J., Park, T., Churchill, M. & Goswami, A. 2020 Wonky whales: the evolution of cranial asymmetry in cetaceans. BMC Biol. 18, 86. (doi:10.1186/s12915-020-00805-4).

[3] Churchill, M., Martinez-Caceres, M., de Muizon, C., Mnieckowski, J. & Geisler, J.H. 2016 The origin of high-frequency hearing in whales. Curr. Biol. 26, 2144–2149. (doi:10.1016/j.cub.2016.06.004).

[4] Galatius, A., Olsen, M.T., Steeman, M.E., Racicot, R.A., Bradshaw, C.D., Kyhn, L.A. & Miller, L.A. 2019 Raising your voice: evolution of narrow-band high-frequency signals in toothed whales (Odontoceti). Biol. J. Linn. Soc. 126, 213–224. (doi:10.1093/biolinnean/bly194).

[5] Arbour, J.H., Curtis, A.A. & Santana, S.E. 2021 Sensory adaptations reshaped intrinsic factors underlying morphological diversification in bats. BMC Biol. 19, 88. (doi:10.1186/s12915-021-01022-3).

[6] Chai, S., Tian, R., Rong, X., Li, G., Chen, B., Ren, W., Xu, S. & Yang, G. 2020 Evidence of echolocation in the common shrew from molecular convergence with other echolocating mammals. Zool. Stud. 59, e4. (doi:10.6620/ZS.2020.59-4).

[7] Galatius, A., Racicot, R., McGowen, M. & Olsen, M.T. 2020 Evolution and diversification of delphinid skull shapes. iScience 23, 101543. (doi:10.1016/j.isci.2020.101543).

[8] Churchill, M., Miguel, J., Beatty, B.L., Goswami, A. & Geisler, J.H. 2019 Asymmetry drives modularity of the skull in the common dolphin (*Delphinus delphis*). Biol. J. Linn. Soc. 126, 225–239. (doi:10.1093/biolinnean/bly190).

[9] Fahlke, J.M., Gingerich, P.D., Welsh, R.C. & Wood, A.R. 2011 Cranial asymmetry in eocene archaeocete whales and the evolution of directional hearing in water. Proc. Natl. Acad. Sci. U.S.A. 108, 14545–14548. (doi:10.1073/pnas.1108927108).

[10] Gatesy, J., Geisler, J.H., Chang, J., Buell, C., Berta, A., Meredith, R.W., Springer, M.S. & McGowen, M.R. 2013 A phylogenetic blueprint for a modern whale. Mol. Phylogen. Evol. 66, 479–506. (doi:10.1016/j.ympev.2012.10.012).

[11] Fahlke, J.M. & Hampe, O. 2015 Cranial symmetry in baleen whales (Cetacea, Mysticeti) and the occurrence of cranial asymmetry throughout cetacean evolution. Sci. Nat. 102, 58. (doi:10.1007/s00114-015-1309-0).

[12] Ness, A.R. 1967 A measure of asymmetry of the skulls of odontocete whales. J. Zool. 153, 209–221. (doi:10.1111/j.1469-7998.1967.tb04060.x).

[13] Huggenberger, S., Leidenberger, S. & Oelschläger, H.H.A. 2017 Asymmetry of the nasofacial skull in toothed whales (Odontoceti). J. Zool. 302, 15–23. (doi:10.1111/jzo.12425).

[14] Cranford, T.W., Amundin, M. & Norris, K.S. 1996 Functional morphology and homology in the odontocete nasal complex: Implications for sound generation. J. Morphol. 228, 223–285.

[15] Mead, J.G. 1975 Anatomy of the external nasal passage and facial complex in the Delphinidae. Washington, D.C., Smithsonian institution press; 72 p.

[16] Thewissen, J.G.M., Cooper, L.N. & Behringer, R.R. 2012 Developmental biology enriches paleontology. J. Vert. Paleontol. 32, 1223–1234. (doi:10.1080/02724634.2012.707717).

[17] Lanzetti, A. 2019 Prenatal developmental sequence of the skull of minke whales and its implications for the evolution of mysticetes and the teeth-to-baleen transition. J. Anat. 235, 725–748. (doi:10.1111/joa.13029).

[18] Roston, R.A. & Roth, V.L. 2019 Cetacean skull telescoping brings evolution of cranial sutures into focus. Anat. Rec. 302, 1055–1073. (doi:10.1002/ar.24079).

[19] Collyer, M.L. & Adams, D.C. 2013 Phenotypic trajectory analysis: Comparison of shape change patterns in evolution and ecology. Hystrix 24, 75–83. (doi:10.4404/hystrix-24.1-6298).

[20] Alberch, P., Gould, S.J., Oster, G.F. & Wake, D.B. 1979 Size and shape in ontogeny and phylogeny. Paleobiology 5, 296–317.

[21] Morris, Z.S., Vliet, K.A., Abzhanov, A. & Pierce, S.E. 2019 Heterochronic shifts and conserved embryonic shape underlie crocodylian craniofacial disparity and convergence. Proc. R. Soc. B 286, 20182389. (doi:10.1098/rspb.2018.2389).

[22] Mitteroecker, P. & Bookstein, F. 2009 The ontogenetic trajectory of the phenotypic covariance matrix, with examples from craniofacial shape in rats and humans. Evolution 63, 727–737. (doi:10.1111/J.1558-5646.2008.00587.X).

[23] Morris, Z.S., Vliet, K.A., Abzhanov, A. & Pierce, S.E. 2021 Developmental origins of the crocodylian skull table and platyrostral face. Anat. Rec. (doi:10.1002/ar.24802).

[24] McGowen, M.R., Tsagkogeorga, G., Álvarez-Carretero, S., dos Reis, M., Struebig, M., Deaville, R., Jepson, P.D., Jarman, S., Polanowski, A., Morin, P.A., et al. 2020 Phylogenomic resolution of the cetacean tree of life using target sequence capture. Syst. Biol. 69, 479–501. (doi:10.1093/sysbio/syz068).

[25] Olson, P.A. 2009 Pilot whales (*Globicephala melas* and *G. macrorhynchus*). In Encyclopedia of Marine Mammals (eds. W.F. Perrin, B. Wursig & J.G.M. Thewissen), pp. 847–852, 2 ed. San Diego, CA, Academic Press.

[26] O’Corry-Crowe, G.M. 2009 Beluga Whale (*Delphinapterus leucas*). In Encyclopedia of Marine Mammals (eds. W. Perrin, B. Wursig & J.H.J. Thewissen), pp. 108–112, 2 ed. San Diego, CA, Academic Press.

[27] Pauly, D., Trites, A.W., Capuli, E. & Christensen, V. 1998 Diet composition and trophic levels of marine mammals. J. Mar. Sci. Technol. 55, 467–481.

[28] Werth, A.J. 2006 Odontocete suction feeding: experimental analysis of water flow and head shape. J. Morphol. 267, 1415–1428. (doi:10.1002/jmor).

[29] McCurry, M.R., Fitzgerald, E.M.G., Evans, A.R., Adams, J.W. & McHenry, C.R. 2017 Skull shape reflects prey size niche in toothed whales. Biol. J. Linn. Soc. 121, 936–946. (doi:10.1093/biolinnean/blx032).

[30] Racicot, R.A. & Preucil, V.E. 2021 Bony labyrinths of the Blackfish (Delphinidae: Globicephalinae). Mar. Mamm. Sci. 38, 29–41. (doi:10.1111/mms.12842).

[31] Jensen, F.H., Johnson, M., Ladegaard, M., Wisniewska, D.M. & Madsen, P.T. 2018 Narrow acoustic field of view drives frequency scaling in toothed whale biosonar. Curr. Biol. 28, 3878–3885. (doi:10.1016/j.cub.2018.10.037).

[32] Racicot, R.A., Darroch, S.A.F. & Kohno, N. 2018 Neuroanatomy and inner ear labyrinths of the narwhal, *Monodon monoceros*, and beluga, *Delphinapterus leucas* (Cetacea: Monodontidae). J. Anat. 223, 421–439. (doi:10.1111/joa.12862).

[33] Turl, C.W. & Penner, R.H. 1989 Differences in echolocation click patterns of the beluga (*Delphinapterus leucas*) and the bottlenose dolphin (*Tursiops truncatus*). J. Acoust. Soc. Am. 86, 497–502. (doi:10.1121/1.398229).

[34] Aguilar Soto, N., Johnson, M.P., Madsen, P.T., Díaz, F., Domínguez, I., Brito, A. & Tyack, P. 2008 Cheetahs of the deep sea: Deep foraging sprints in short-finned pilot whales off Tenerife (Canary Islands). J. Anim. Ecol. 77, 936–947. (doi:10.1111/j.1365-2656.2008.01393.x).

[35] May-Collado, L.J., Agnarsson, I. & Wartzok, D. 2007 Phylogenetic review of tonal sound production in whales in relation to sociality. BMC Evol. Biol. 7, 136. (doi:10.1186/1471-2148-7-136).

[36] Navalón, G., Nebreda, S.M., Bright, J.A., Fabbri, M., Benson, R.B.J., Bhullar, B.-A., Marugán-Lobón, J. & Rayfield, E.J. 2021 Craniofacial development illuminates the evolution of nightbirds (Strisores). Proc. R. Soc. B 288, 20210181. (doi:10.1098/rspb.2021.0181).

[37] Frainer, G., Moreno, I.B., Serpa, N., Galatius, A., Wiedermann, D. & Huggenberger, S. 2019 Ontogeny and evolution of the sound-generating structures in the infraorder Delphinida (Odontoceti: Delphinida). Biol. J. Linn. Soc. 128, 700–724. (doi:10.1093/biolinnean/blz118/5550896).

[38] Galatius, A. 2010 Paedomorphosis in two small species of toothed whales (Odontoceti): how and why? Biol. J. Linn. Soc. 99, 278–295. (doi:10.1111/j.1095-8312.2009.01357.x).

[39] Frainer, G., Huggenberger, S., Moreno, I.B., Plön, S. & Galatius, A. 2021 Head adaptation for sound production and feeding strategy in dolphins (Odontoceti: Delphinida). J. Anat. 238, 1070–1081. (doi:10.1111/JOA.13364).

[40] Berta, A. & Lanzetti, A. 2020 Feeding in marine mammals: an integration of evolution and ecology through time. Palaeontol. Electron. 23, a40. (doi:doi.org/10.26879/951).

[41] Frainer, G., Huggenberger, S. & Moreno, I.B. 2015 Postnatal development of franciscana’s (*Pontoporia blainvillei*) biosonar relevant structures with potential implications for function, life history, and bycatch. Mar. Mamm. Sci. 31, 1193–1212. (doi:10.1111/MMS.12211).

[42] Galatius, A. & Goodall, R.N.P. 2016 Skull shapes of the Lissodelphininae: Radiation, adaptation and asymmetry. J. Morphol. 277, 776–785. (doi:10.1002/jmor.20535).

[43] Churchill, M., Geisler, J.H., Beatty, B.L. & Goswami, A. 2018 Evolution of cranial telescoping in echolocating whales (Cetacea: Odontoceti). Evolution 72, 1092–1108. (doi:10.1111/evo.13480).

[44] Kuzmin, A.A. 1976 Embryogenesis of the osseus skull of the sperm whale. In Investigations on Cetacea (ed. G. Pilleri), pp. 187–202. Berne, Switzerland, Institute of Brain Anatomy, University of Berne.

[45] Lambert, O. 2005 Phylogenetic affinities of the long-snouted dolphin *Eurhinodelphis* (Cetacea, Odontoceti) from the Miocene of Antwerp, Belgium. Palaeontology 48, 653–679. (doi:10.1111/j.1475-4983.2005.00472.x).

[46] Geisler, J.H., McGowen, M.R., Yang, G. & Gatesy, J. 2011 A supermatrix analysis of genomic, morphological, and paleontological data from crown Cetacea. BMC Evol. Biol. 11, 112. (doi:10.1186/1471-2148-11-112).

[47] Murakami, M., Shimada, C., Hikida, Y., Soeda, Y. & Hirano, H. 2014 *Eodelphis kabatensis*, a new name for the oldest true dolphin *Stenella kabatensis* Horikawa, 1977 (Cetacea, Odontoceti, Delphinidae), from the upper Miocene of Japan, and the phylogeny and paleobiogeography of Delphinoidea. J. Vert. Paleontol. 34, 491–511. (doi:10.1080/02724634.2013.816720).

[48] Galatius, A., Berta, A., Frandsen, M.S. & Goodall, R.N.P. 2011 Interspecific variation of ontogeny and skull shape among porpoises (Phocoenidae). J. Morphol. 272, 136–148. (doi:10.1002/jmor.10900).

[49] Cooper, L.N., Sears, K.E., Armfield, B.A., Kala, B., Hubler, M. & Thewissen, J.G.M. 2018 Review and experimental evaluation of the embryonic development and evolutionary history of flipper development and hyperphalangy in dolphins (Cetacea: Mammalia). Genesis 56, e23076. (doi:10.1002/dvg.23076).

[50] Armfield, B.A., Zheng, Z., Bajpai, S., Vinyard, C.J. & Thewissen, J.G.M. 2013 Development and evolution of the unique cetacean dentition. PeerJ 1, e24. (doi:10.7717/peerj.24).

[51] Lanzetti, A., Chrouch, N., Miguez, R.P., Fernandez, V. & Goswami, A. 2022 Developing echolocation: distinctive patterns in the ontogeny of the tympanoperiotic complex in baleen and toothed whales (Cetacea). Biol. J. Linn. Soc. 135, 394–406. (doi:10.1093/biolinnean/blab160).

[52] R Core Team. 2021 R: A language and environment for statistical computing. See http://www.r-project.org/.

[53] Baken, E.K., Collyer, M.L., Kaliontzopoulou, A. & Adams, D.C. 2021 geomorph v4.0 and gmShiny: enhanced analytics and a new graphical interface for a comprehensive morphometric experience. Methods Ecol. Evol. 12, 2355–2363. (doi:10.1111/2041-210x.13723).

[54] Guillerme, T., Weisbecker, V. & Marcy, A.E. 2019 landvR: Tools for measuring landmark position variation. See https://doi.org/10.5281/zenodo.2620785.

[55] Mead, J.G. & Fordyce, R.E. 2009 The therian skull: a lexicon with emphasis on the odontocetes. Washington, D.C., Smithsonian institution press; 249 p.

[56] Lindeløv, J.K. 2020 mcp: An R package for regression with multiple change points. OSF Preprints. (doi:10.31219/osf.io/fzqxv).

[57] Collyer, M.L. & Adams, D.C. 2021 RRPP: Linear model evaluation with randomized residuals in a permutation procedure. See https://cran.r-project.org/web/packages/RRPP.

[58] Revell, L.J. 2012 phytools: An R package for phylogenetic comparative biology (and other things). Methods Ecol. Evol. 3, 217–223. (doi:10.1111/j.2041-210X.2011.00169.x).

[59] Lenth, R.V. 2022 emmeans: estimated marginal means, aka least-squares mean. See https://cran.r-project.org/web/packages/emmeans.

[60] Wickham, H. 2016 ggplot2: Elegant graphics for data analysis. See https://ggplot2.tidyverse.org.

