## Supplementary material for "The ontogeny of asymmetry in echolocating whales": electronic supplementary material

**This PDF file includes:**

Supplemental results
Supplemental methods
Figures S1 to S12
Tables S1 to S8
Legends for Datasets S1 to S2
Supplemental references

**Other supplementary materials for this manuscript include the following:**

Datasets S1 to S2

### Supplemental results

#### (a) Principal Components Analysis (PCA) of skull shape

##### (i) Raw data

A PCA of GPA-aligned skull shape data demonstrates a high degree of morphological convergence between *Delphinapterus* and *Globicephala* (electronic supplementary material, figure S7*a*, table S6). Similarity in their skull shape is recognizable in early ontogenetic stages, which cluster on the negative size of PC1 (56.95% of total variation), and the two taxa continue to occupy the same area of the morphospace in the postnatal stages. Along PC1, there is an increase in size and asymmetry level correlated with maturity, with the more asymmetrical adults occupying the positive side of this axis. The progressive posterior shift and overlap of the skull bones, known as telescoping [1], and rostral elongation are the most notable changes in skull shape that occur on this axis and characterize ontogenetic growth. PC2 (14.53%) instead divides the dataset phylogenetically and by morphotype (electronic supplementary material, figure S7*a*). *Delphinapterus* and *Globicephala*, which share a broader rostrum indicative of suction feeding [2], fall on the negative end, and the other taxa with more elongated rostra at the positive one. There is also a more subtle size and asymmetry gradient on this axis, with larger and more asymmetric specimens at the negative end. *Phocoena* occupies an intermediate area closer to *Stenella* and *Lagenorhynchus*, owing to their comparable size and relative rostral elongation, but its range does not similarly expand to the extremes of PC1, probably due to its lower level of asymmetry.

##### (ii) Size-corrected

A PCA on the shape residuals of the common allometric component further clarifies these patterns of morphological convergence (electronic supplementary material, figure S7*b*, table S6). PC1 (43.1%) separates specimens by taxon, but also clearly exemplifies differences in rostral morphology. *Delphinapterus* and *Globicephala* with their broader and shorter rostra occupy an overlapping region at the negative end of the axis, while the longirostrine *Lagenorhynchus* and *Stenella* are at the opposite side. *Phocoena* remains in the middle, but it is well separated from the other taxa by its position on PC2 (12.89%). This taxon occupies the positive side of this axis due to its low level of asymmetry, while the more asymmetric adult specimens of the other taxa are at the opposite end.

### Supplemental methods

#### (a) Dataset composition

Some of the genera used in the study are monospecific (i.e. *Delphinapterus*) while others are comprised of multiple species and might be paraphyletic (i.e. *Lagenorhynchus* [3]). Though, all the species in each genus are similar morphologically and ecologically, with molecular data being necessary to identify their precise taxonomy in some cases [3, 4], making it difficult to confidently assign museum specimens precisely to a species when precise information might be lacking, and the original species name used is now considered invalid. But these taxa are distantly related from each other and well separated phylogenetically [4]. Therefore, specimens from any species in that genus can be pulled in one taxonomic unit to be used in the analyses.

Approximate age of the specimens was determined based on total length of the specimen.

This was either measured directly for specimens in fluid collections, or it was extracted from the literature or collection metadata. If no information on total length was available, it was estimated based on skull measurements, which is a reliable method to reconstruct body size in Cetacea [5]. For prenatal specimens, the approximate age as a proportion of gestation length was reconstructed using growth curves and information from the literature for each taxon [6-9] following the methodology used in [10]. The gestation was binned into two growth stages for analyses: early fetus (from embryonic stages to 50% of gestation time) and late fetus and neonate (from 50% gestation time to birth, including the first year of life). This was done to ensure that small errors in the assessment of the age of the specimens would not undermine the results, as for most species basic information such as the length at birth is unknown or contrasting information have been published in the literature [6, 9]. Postnatal development was divided into two stages: juvenile and adult. Based on their total length, specimens were determined juveniles if their length was less than the estimated size at sexual maturity for that species according to published data [11-17].

#### (b) Shape data acquisition and rendering

Specimens were digitized specifically for this study from the Natural History Museum, London (NHMUK) collections. CT data and surface scans of specimens from other institutions were also used with permission (electronic supplementary material, dataset 1).

X-ray computed tomography (CT) datasets presented in this study originates from two institutes: The Imaging and Analysis Centre of the Natural History Museum (London, UK) and the Bio-Imaging Research Center of the Smithsonian Institution (Washington, D.C., USA). All parameters and instruments are listed in Dataset S2. For specimens imaged at the NHMUK, parameters were adjusted to maximise resolution, and ensure a decent X-ray transmission and signal-to-noise ratio. For specimen larger than the field of view, several CT acquisitions were performed along the vertical axis. These datasets were subsequently merged using the protocol described in [18] and are indicated as ‘vertical stitching’ in the column ‘X-ray CT modality (electronic supplementary material, dataset 2).

Surface scans were performed at 0.2-0.5 mm resolution depending on the size of the specimen.

All CT data were first imported in ImageJ [19] and scaled down by a factor of 2 in all 3 dimensions (i.e., binning 2x2x2), reducing the image size and limiting the number of slices to a maximum of 1500 to aid with segmentation, and also cropped and adjusted for brightness/contrast where needed. Images were then exported as 16-bit TIFF images

stacks for segmentation in Avizo Lite v. 2020.3. Skull bones were manually segmented and exported as OBJ surface files. Skulls of disarticulated osteological specimens were digitally reconstructed by segmenting bone element. For surface scans, the dorsal and ventral surface of the skull digitized separately were merged in VXElements v. 8.1.
All meshes were imported in Geomagic Wrap 2017 for surface processing. Disarticulated osteological specimens were reconstructed aligning them to a model of a skull of similar length and species. In all skull meshes, holes were filled, then in order to obtain a consistent smooth surface and eliminate small defects that might influence the results, spikes were removed, and a quick smooth was performed. All surfaces were sampled to 1,500,000 triangles to ensure a consistent visualization for landmarking and then exported as PLY files.

**(c) Landmarking and data preparation**
The 64 Type I and Type II single points landmarks [20], and associated semilandmarks curves, were selected to represent the full shape of the cranium while being identifiable from the earliest stages of growth. Unidentifiable landmarks and curves, either due to broken parts of the specimen or due to the absence of that bone at that developmental stage, were marked as missing during landmarking. We placed 'missing' landmarks as close to the missing structure as possible and marked it as a 'missing landmark' in Stratovan Checkpoint v. 2020.10.13.0859, which automatically assigns a coordinate of −9999. 24 of the 58 specimens had at least one fixed landmark or curved marked as missing. Of these, 14 had "absent" structures (i.e., nasal bones not ossified yet in early fetus stage, interparietal not visible in adults). Therefore, only 17% of the samples as some broken feature to be estimated before performing GM analyses.
After the exported landmark coordinates were loaded into R v. 4.1.1 [21] for analyses, curves were resampled to set the same number of semilandmark points for each for all specimens using the package SURGE [22], and then slid on the surface using a custom function to minimize bending energy and ensure homology [23].
We then used the 'fixLMtps' function from the R package 'Morpho' v. 2.9 [24, 25] to estimate missing landmarks by mapping weighted averages from three similar, complete configurations onto the missing specimen. Estimated landmarks are then added to the deficient configuration [25]. The deformation is performed by a thin-plate-spline interpolation calculated using the available landmarks [20, 25]. For genuinely absent bones, all landmarks and semilandmarks associated with that bone were placed in a single "zero-area" point adjacent to its position in other taxa, following the method described by [23]. List of specimens with fixed LMs or curves marked as absent is available on GitHub at https://github.com/AgneseLan/ontogeny-asymmetry-dolphin (https://doi.org/10.5281/zenodo.6553285).

**(d) Data analyses**
**(i) Asymmetry analysis**
First, we assessed if directional asymmetry (DA) was present in the entire dataset, in each taxon and in each of the four growth categories with the 'bilat.symmetry' function in 'geomorph' [26]. Having noticed the expected patterns based on the literature [27, 28], we then proceeded to calculate DA shapes and symmetric shape for each specimen using the function iteratively on one specimen at a time. The function produces two full DA shapes configurations per specimen, one representing the asymmetric component of the left side and its mirror on the right side to recreate a full configuration, and the other the asymmetric component of the right side mirrored on the left. The 'bilat.symmetry' functions also produces a asymmetric shape component, but we did not it for the analyses as that is also influenced by fluctuating asymmetry (FA), which we did not want

to consider in the study. The symmetric and DA shapes for each specimen calculated iteratively using ‘bilat.asymmetry’ were combined in a single array and Procrustes superimposition was performed again to ensure that all shapes and specimens were aligned correctly (‘gpagen’ function). The Euclidean distance between the landmarks of the first part of the array representing symmetric shape and the ones in the second part of the array presenting DA shape was calculated with the ‘linear.dist’ function in ‘landVR’ [29]. Observing the distribution of the distance variations in each phase of growth, we noticed that most of the asymmetry is due to the progressive shifts in interparietal bone position (electronic supplementary material, figure S9). This bone is not visible or is greatly reduced in the adults of all taxa analysed except *Phocoena*, as it fuses and folds underneath the supraoccipital after birth as part of the telescoping process [1, 30]. If not visible in a specimen, the landmarks placed on the anterior end of the interparietal were therefore marked as “absent” in the GM analysis with their position assigned arbitrarily. To avoid the asymmetry analysis to be skewed by this, we run the same analyses starting from the GPA excluding the three landmarks placed on the interparietal. The patterns of ontogenetic change in asymmetry and the differences among taxa and ontogenetic stages are broadly similar between the two analyses, whole skull (electronic supplementary material, figure S9-S10-S11, table S8) and interparietal excluded (figure 1, electronic supplementary material, figure S1-S2, table S1), and therefore we decided to discuss in detail the results of the configuration with the interparietal removed, allowing us to better visualize variation in symmetry in other parts of the neurocranium. After calculating the linear distances between DA and symmetric shapes using ‘landVR’, we obtained a matrix of 64x58 distances for the whole skull and 61x58 when excluding the three landmarks on the interparietal (n. 57-58-59), one for each fixed landmark point for each specimen. While the left and right DA configurations are different from each other given that DA is significant in the dataset, we found no difference in the mean linear distances between each of the DA shapes and the symmetric shapes for the entire dataset. Therefore, we used the right-side DA shape component for all analyses. The distances representing the distribution of asymmetry were visualised by applying a colour function to average specimen’s landmark configurations using the ‘rgl’ package v. 0.107.14 [31].

**(ii) Shape variation**

The PCA was performed both on the raw GPA-aligned data (electronic supplementary material, figure S7A) and on the residuals of the common allometric regression (electronic supplementary material, figure S7B), obtained using the ‘procD.lm’ function in geomorph with the log-transformed CS as independent variable and Procrustes shape coordinates as the response variable. Correlation between the first two PC components, representing most of the variation, with size, growth stage, phylogeny as genus, and asymmetry as mean DA to symmetric shape distance were calculated using the ‘lm’ function in the stats package [21] for the raw shape dataset. Only the effect of phylogeny and asymmetry were tested for the residuals’ dataset.

**(iii) Allometry analysis**

To calculate differences between prenatal and postnatal allometry, we used shape regression scores obtained from a ‘procD.lm’ model (‘geomorph’ package [26]) that accounted for changes in slope between genera (shape ~ logCS * genera) as input for the ‘mcp’ package [32] following [33]. We compared: (1) a common model with no break points, (2) a model with varying slopes among taxa but no break points, (3) a model with one common breakpoint for all data, and (4) a model with varying breakpoints and slopes among taxa. The break point was consistently positioned in the prenatal portion of growth, well below the estimated size at birth, pointing to a change in the rate of growth in shape and size during the prenatal period (electronic supplementary material,

figure S12*a*). The model with one break point but common slopes though also highlighted a possible change around birth electronic supplementary material, figure S12*b*). Given the strong support for the one break point model and the fact that the data are binned in growth stages that span the border between prenatal and postanal ontogeny, we decided to perform the analyses considering only one allometric rate shift, with specimens categorized as early fetus and late fetus/neonate to represents prenatal growth and juvenile and adults for postnatal. It is possible that with a more granular binning of growth stages multiple break points might be identified in the ontogeny of toothed whales, but applying this model already allows to better characterize changes in allometry among taxa and the heterochronic process at play than using a single rate of growth for the entire development (electronic supplementary material, table S3). To quantitatively compare ancestral and modern slopes, it was necessary to create predicted regression shape scores and size data based on the extant dataset using base R 'stat' functions. These data were then used along with the regression scores and sizes for the extant genera calculated using 'procD.lm' to create simple regression models ('lm' function) to compare in 'emmeans' [34]. This implies that the quantitative comparisons between ancestral and modern slopes might be unreliable due to the nature of the data used, though the qualitive comparison remain valid and a valuable insight into the evolution of allometry and its effect on the evolution and ontogeny of asymmetry in odontocetes.
The code use for the analyses is available on GitHub at https://github.com/AgneseLan/ontogeny-asymmetry-dolphin (https://doi.org/10.5281/zenodo.6553285).

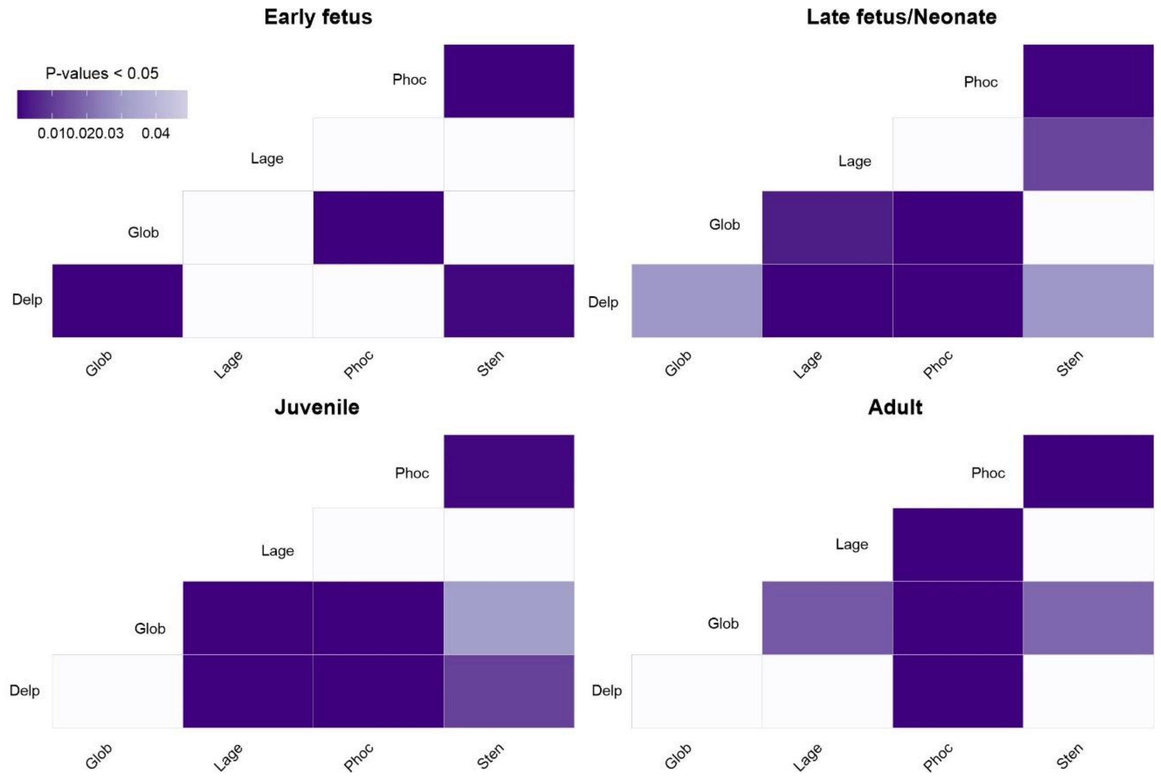


**Figure S1.** Significance patterns of pairwise comparison of mean distances between DA and symmetric shape among taxa for each growth stage, fixed landmarks for interparietal excluded (LM 57-58-59). Delp = *Delphinapterus*, Glob = *Globicephala*, Lage = *Lagenorhynchus*, Phoc = *Phocoena*, Sten = *Stenella*.

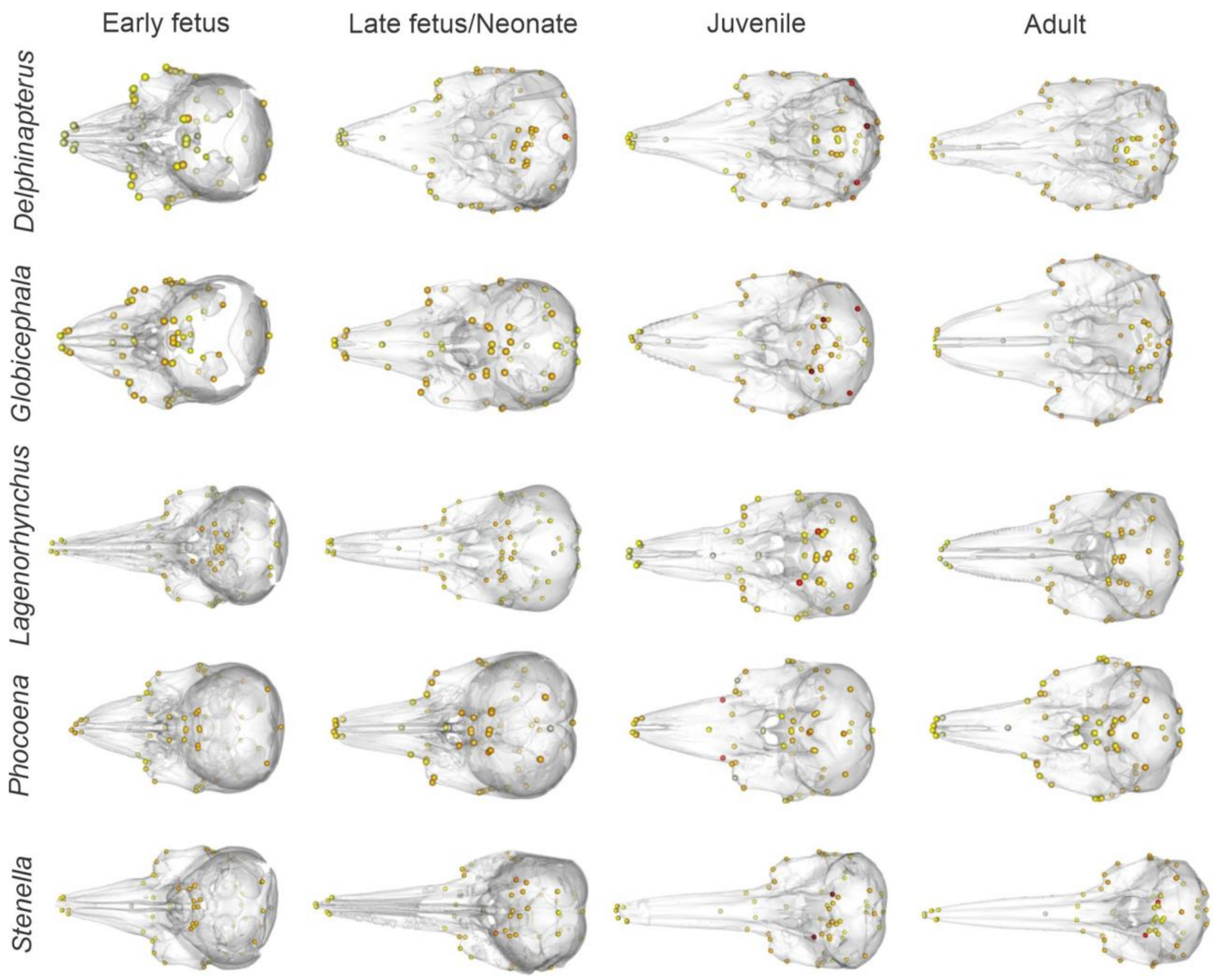


**Figure S2**. Variance of mean distances between DA and symmetric shape for each landmark across taxa and growth stages, fixed landmarks for interparietal excluded (LM 57-58-59). Darker colour represents landmarks with higher distance, or larger deviation from symmetric shape, relative to the mean distances of the entire dataset. Skulls not to scale.

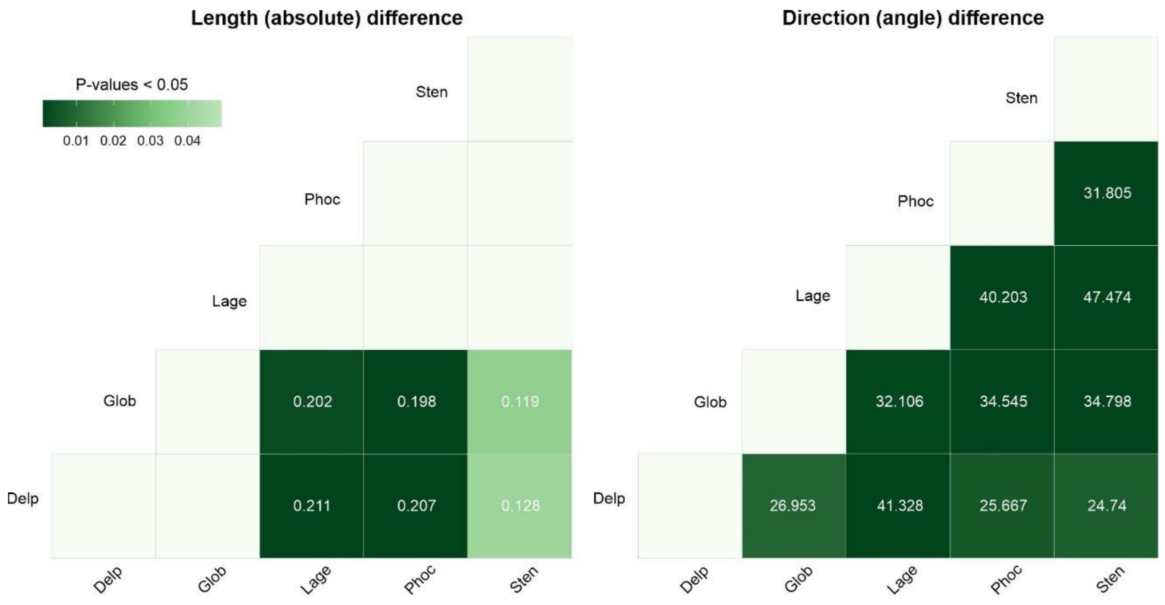


**Figure S3**. Significance patterns of pairwise comparison phenotypic trajectory analysis of skull shape development among genera. “d” values for length and “angle” value for direction are reported for significantly different pairs. Shape metric did not show any significant differences. Delp = *Delphinapterus*, Glob = *Globicephala*, Lage = *Lagenorhynchus*, Phoc = *Phocoena*, Sten = *Stenella*. Full results in table S2.

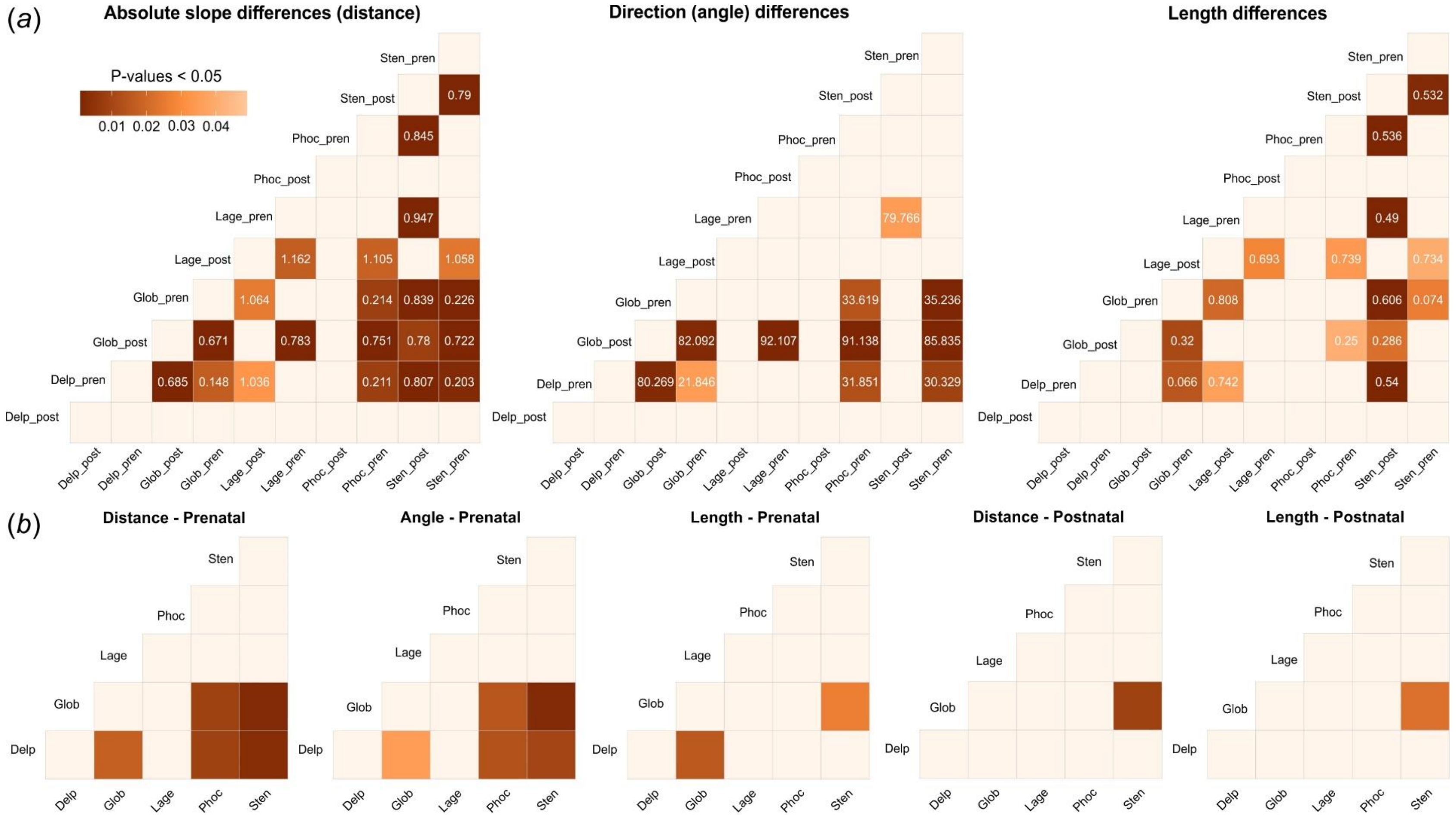


**Figure S4**. Significance patterns of pairwise comparison of allometry slopes among ancestral states and genera. (*a*) All genera and ancestral states and stages; (*b*) All genera and ancestral states divided in prenatal and postnatal stages. 1 = ancestor of Delphinoidea, 2 = ancestor of Monodontidae and Phocoenidae, 3 = ancestor of Delphinidae, 4 = ancestor of *Stenella* and *Globicephala*, Delp = *Delphinapterus*, Glob = *Globicephala*, Lage = *Lagenorhynchus*, Phoc = *Phocoena*, Sten = *Stenella*. Full results in table S4.

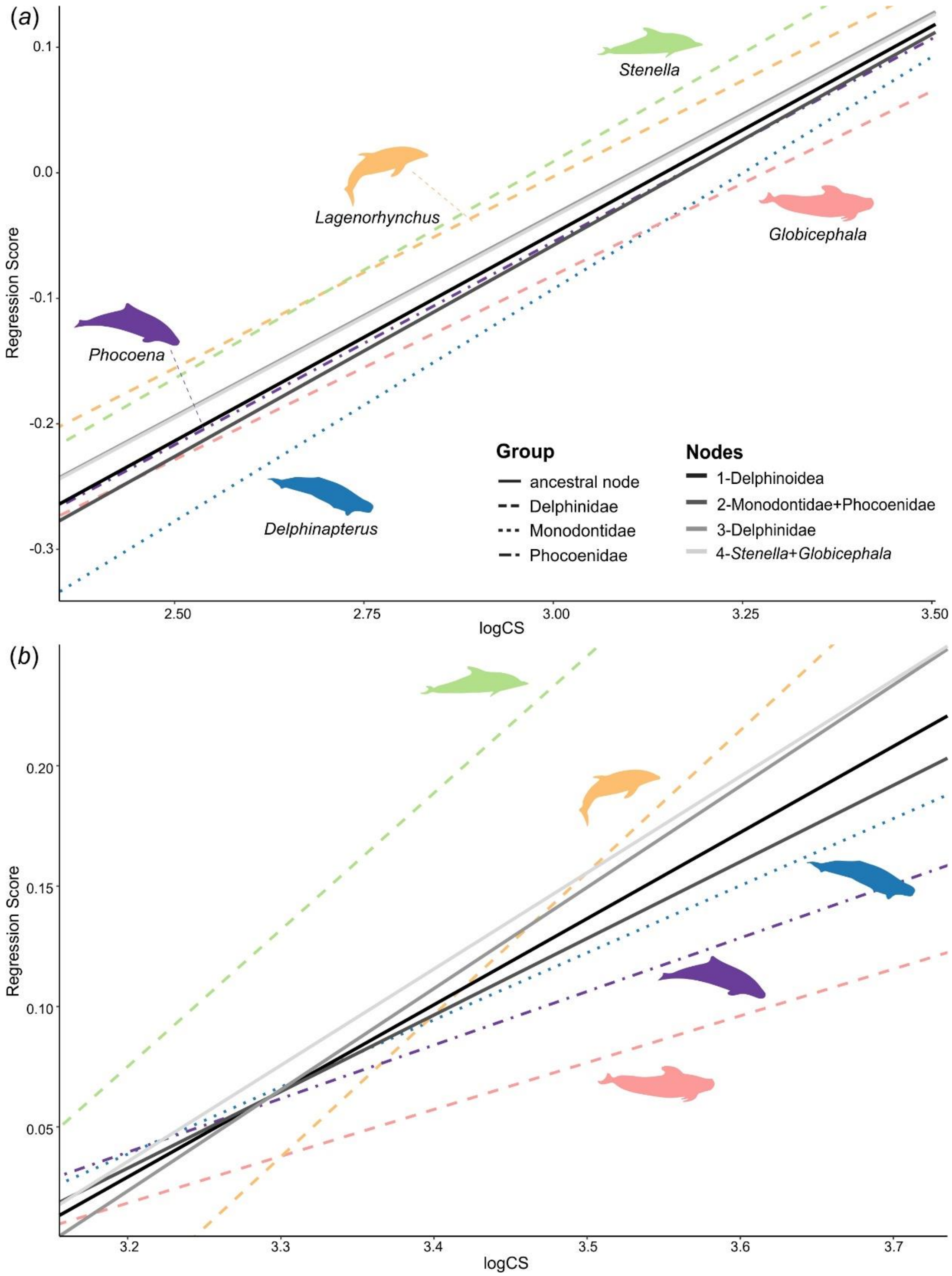


**Figure S5**. Allometric regression of skull shape and size (logCS) for each taxon and ancestral state divided by stage. (*a*) Prenatal allometry; (*b*) Postnatal allometry.

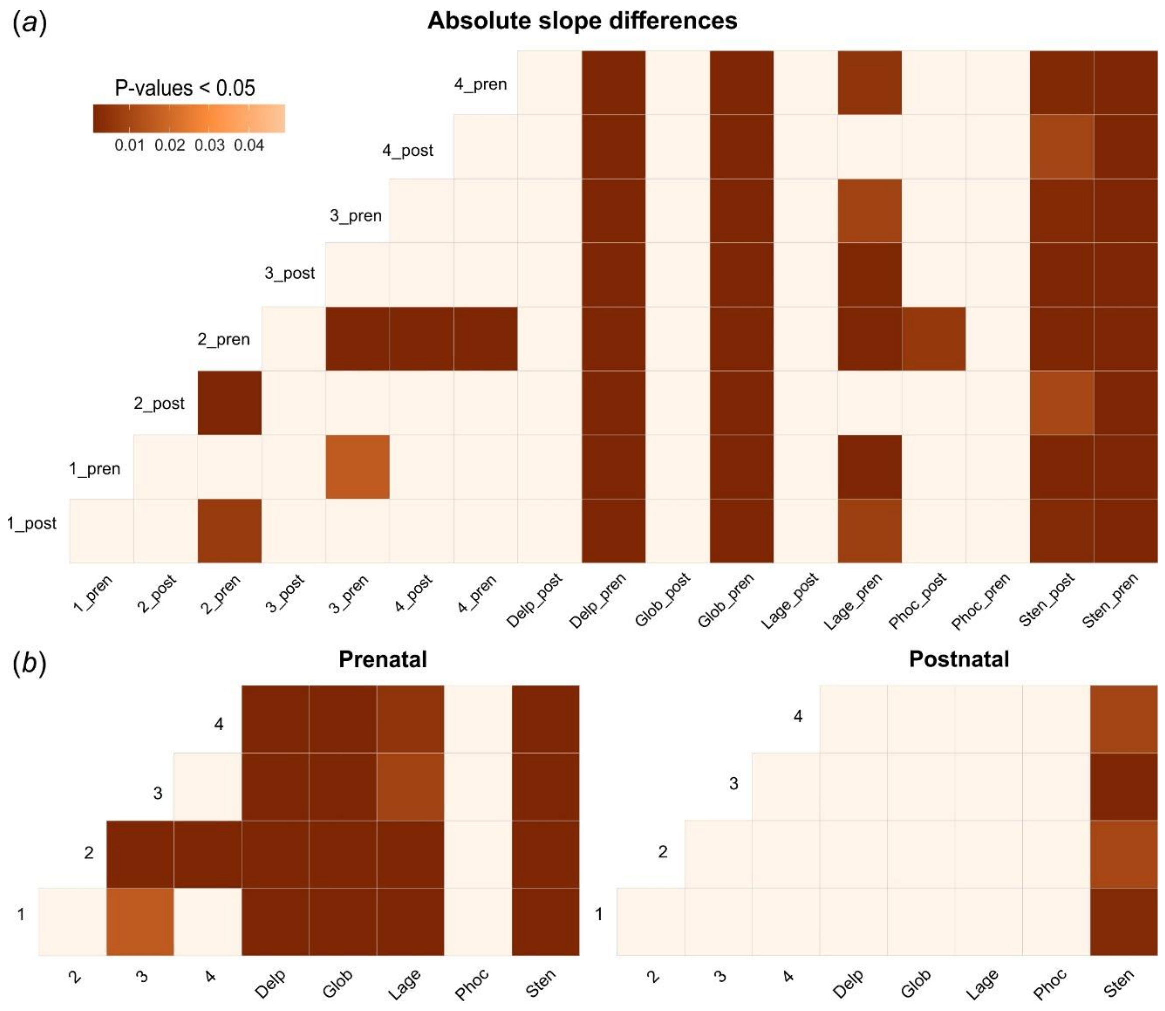


**Figure S6**. Significance patterns of pairwise comparison of allometry slopes among genera. (*a*) All genera and stages; (*b*) All genera divided in prenatal and postnatal stages. “d” values for distance and length and “angle” value for direction are reported for significantly different pairs. Direction metric did not show any significant differences for postnatal stage. Delp = *Delphinapterus*, Glob = *Globicephala*, Lage = *Lagenorhynchus*, Phoc = *Phocoena*, Sten = *Stenella*. Full results in table S5.

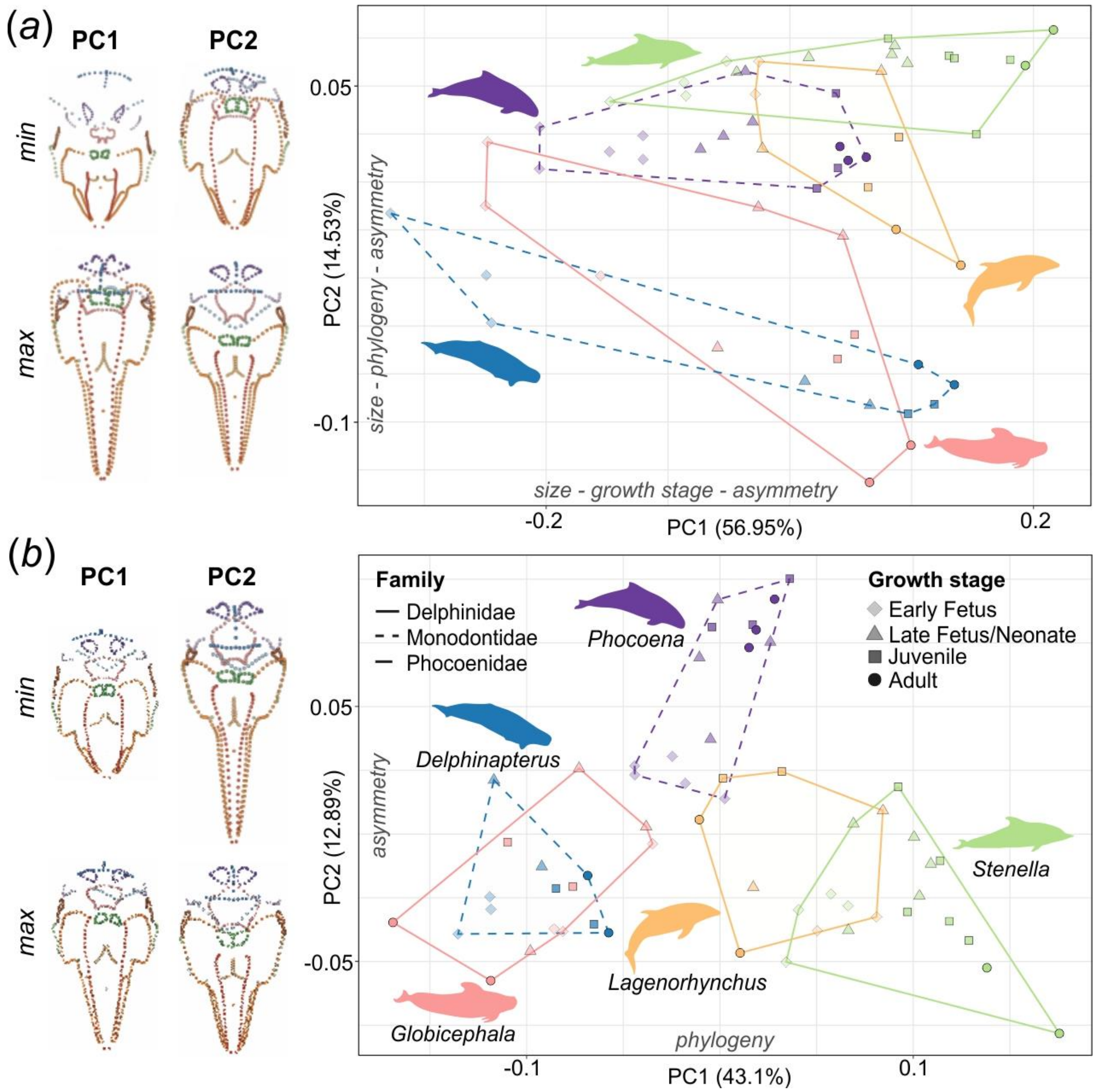


**Figure S7**. Skull ontogeny morphospaces (PCA), showing the shape variation through ontogeny and phylogeny in the dataset, with a significant overlap between Delphinapterus and Globicephala. (*a*) Raw data; (*b*) Size-corrected. The landmark plots on the left represent the morphological extremes on the PC1 and PC2 axis for each plot, in dorsal view. Fixed landmarks and semi-landmarks are represented in unique colours to highlight shape changes in different regions of the skull. The factors that significantly explain shape distribution are reported on each axis. See table S6 for details.

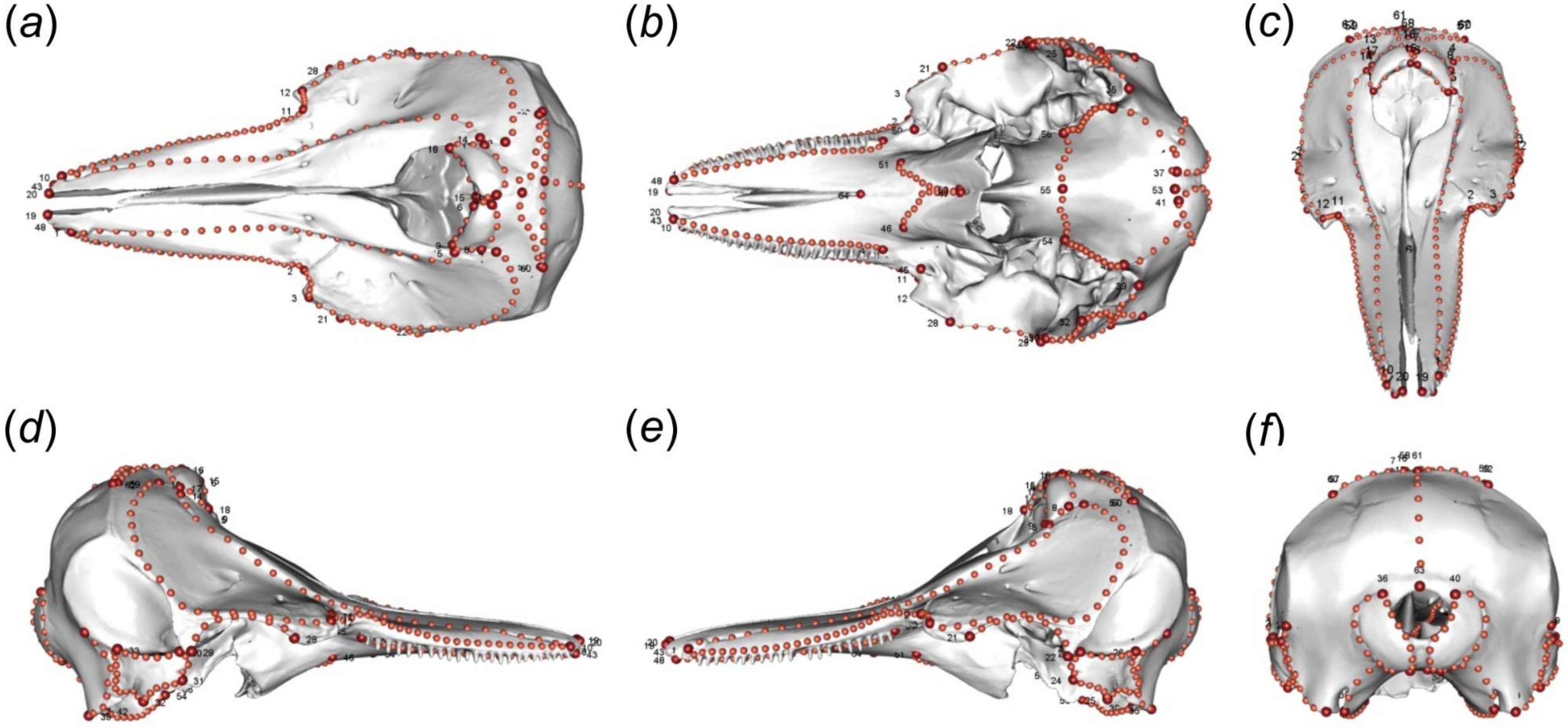


**Figure S8**. Landmarking scheme used in this study, plotted on the skull of an adult *Lagenorhynchus* (*L. albirostris*, AMNH 37162). (*a*) Dorsal view; (*b*) Ventral view; (*c*) Anterior view; (*d*) Medial view; (*e*) Lateral view; (*f*) Posterior view. Fixed landmark points in red and numbered, curve semi-landmarks in orange. See table S7 for details.

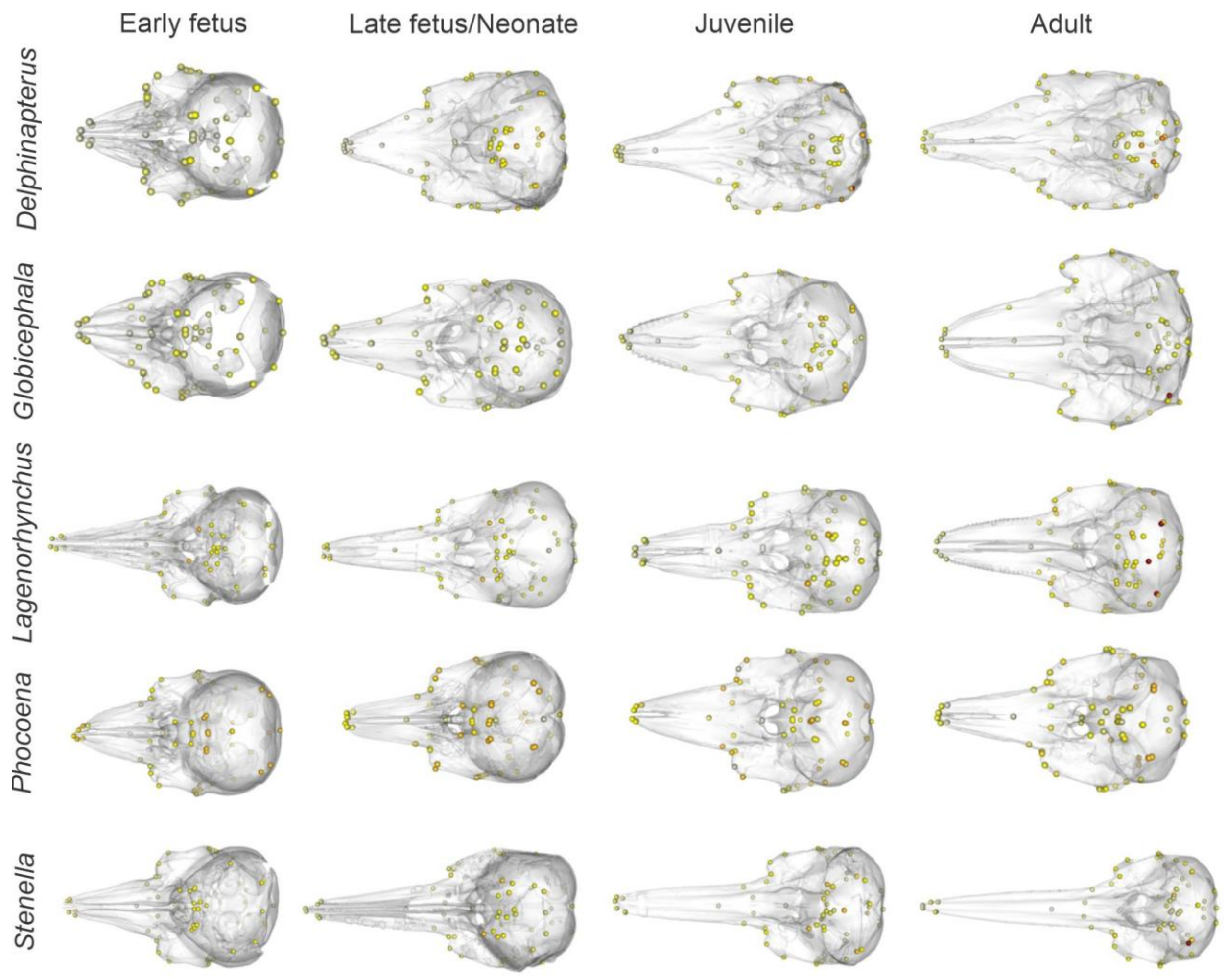


**Figure S9**. Variance of mean distances between DA and symmetric shape for each landmark across taxa and growth stages, all fixed landmarks included. Darker colour represents landmarks with higher distance, or larger deviation from symmetric shape, relative to the mean distances of the entire dataset. Skulls not to scale.

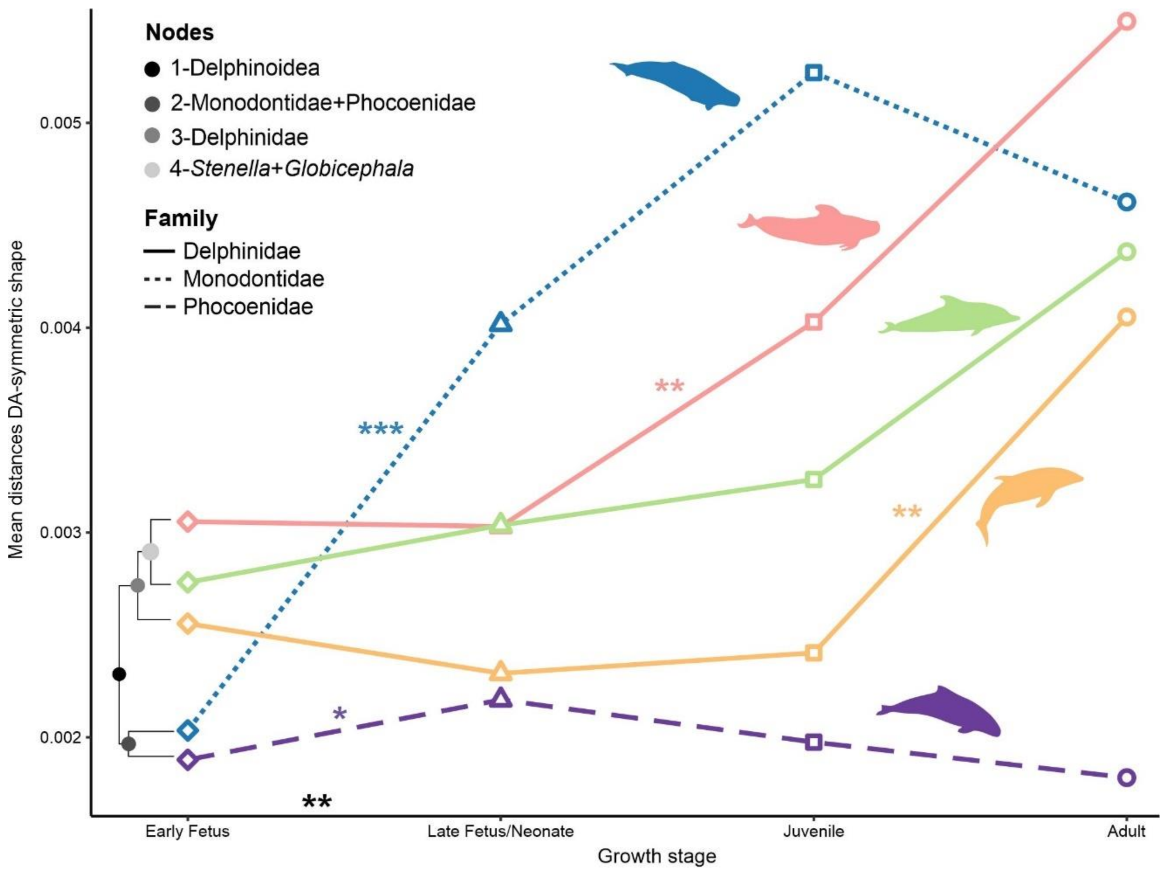


**Figure S10**. Patterns of change in asymmetry though ontogeny of selected taxa, all fixed landmarks included. Degree of asymmetry is quantified as the mean distance between the directional asymmetric (DA) and symmetric shape across all landmarks for all specimens of the same genus at each growth stage. The phylogeny drawn on the left highlights the close correspondence between phylogenetic structure and starting level of asymmetry in ontogeny. Significant shifts in asymmetry levels between growth stages in the entire dataset and per genus are indicated by asterisks (‘***’ p<0.001, ‘**’p<0.01, ‘*’ p<0.05). Full results in table S8 and figure S9-S11.

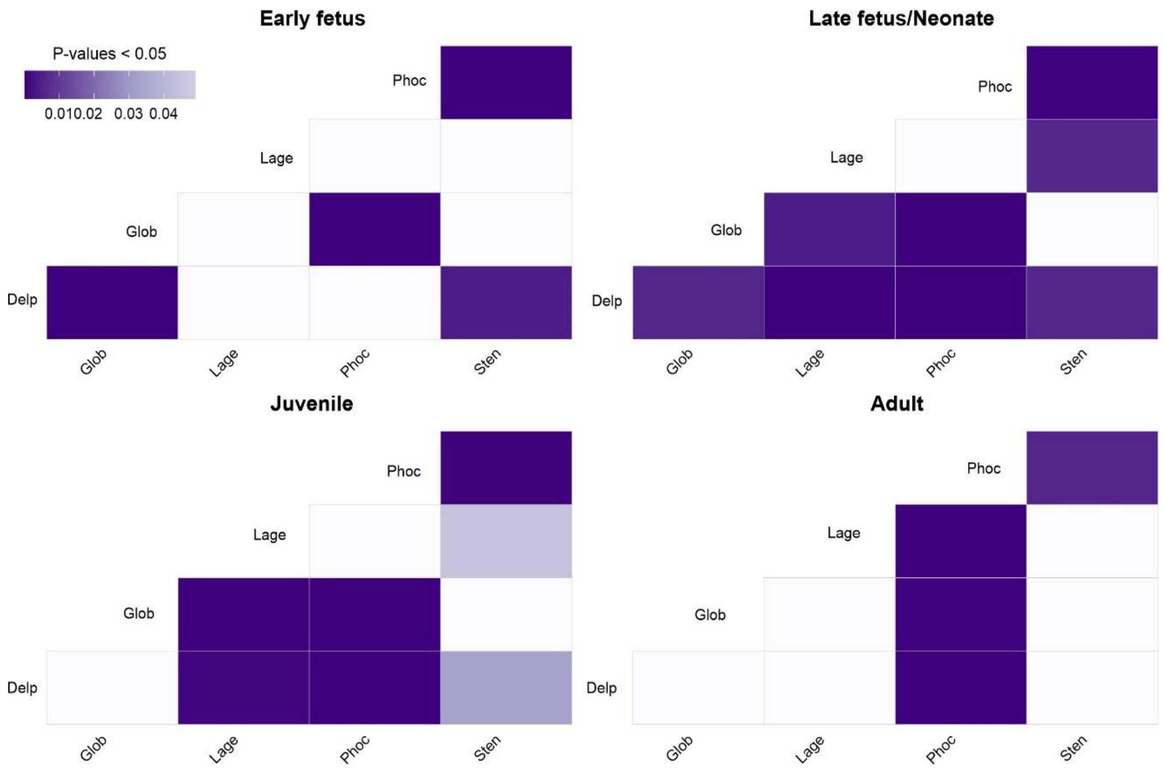


**Figure S11**. Significance patterns of pairwise comparison of mean distances between DA and symmetric shape among taxa for each growth stage, all fixed landmarks included. Delp = *Delphinapterus*, Glob = *Globicephala*, Lage = *Lagenorhynchus*, Phoc = *Phocoena*, Sten = *Stenella*.

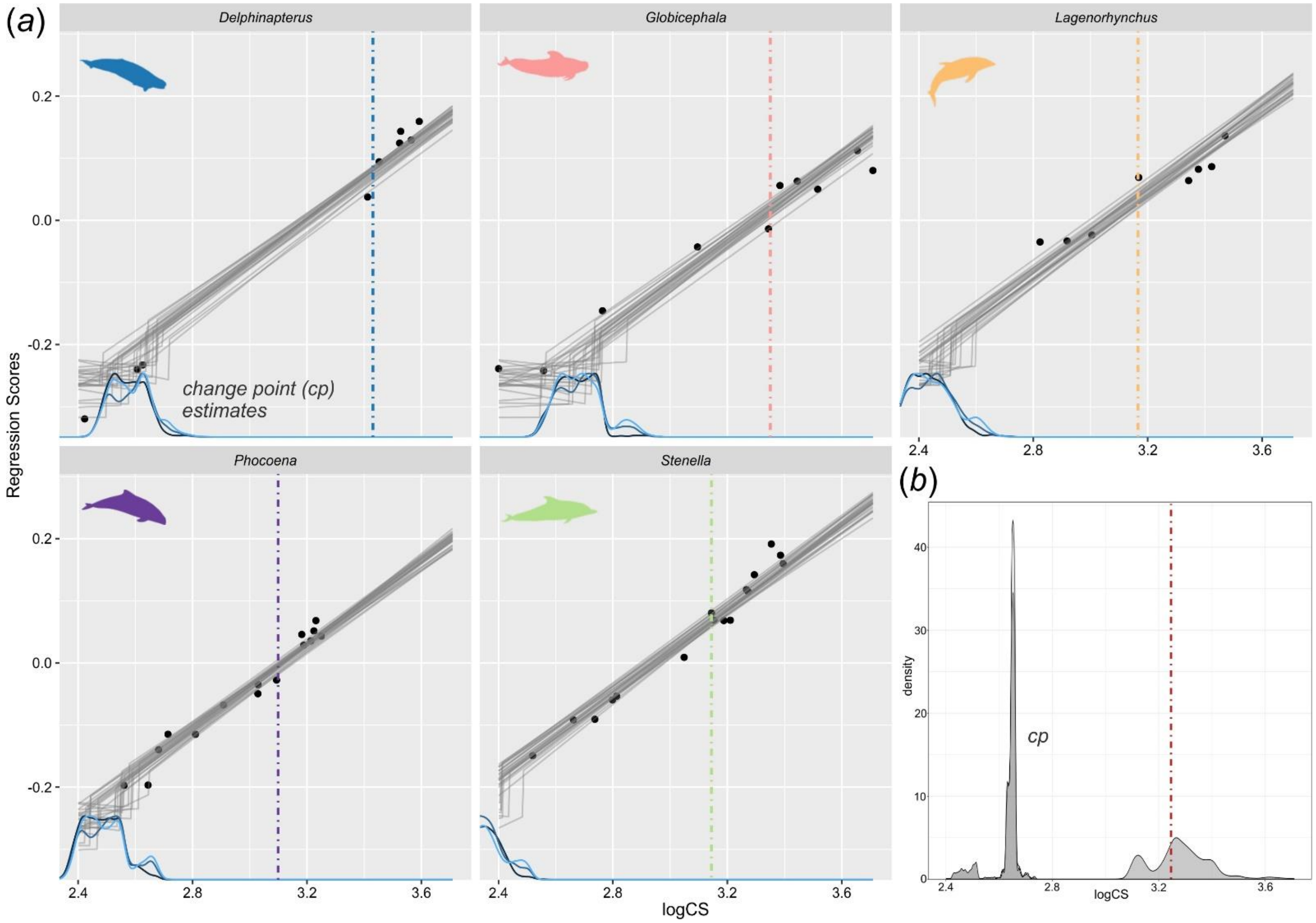


**Figure S12.** Plots of best model and one breakpoint model from multiple breakpoint test for allometry regression. (*a*) Regression plots of regression scores and size (logCS) of best model: one breakpoint varying by taxon with density plot of change points (cp) estimates; (*b*) Density plot of cp estimates for one breakpoint model common to all taxa. Dot-dash line indicates mean size of neonates in the sample for each taxon (*a*) and across all taxa (*b*). See table S3 and supplemental methods for additional details.

**Table S1.** Pairwise comparison of mean distances between DA and symmetric shape among consecutive growth stages for entire dataset and by taxon, fixed landmarks for interparietal excluded (LM 57-58-59). Distance between asymmetric and symmetric shape for each specimen calculated using the 'landVR' package [29] and then analysed grouped by growth stage. Difference between means (t) and p-value obtained with a Welch Two Sample t-test are reported for each pair. Significant p-values (p<0.05) are in bold.

| | Entire dataset | *Delphinapterus* | *Globicephala* | *Lagenorhynchus* | *Phocoena* | *Stenella* |
|---|---|---|---|---|---|---|
| Early fetus – Late fetus/neonate | **-2.9696**<br>**p = 0.0036** | **-7.1427**<br>**p = 2.53E-10** | 0.0128<br>p = 0.9898 | 0.8205<br>p = 0.4140 | **-2.4537**<br>**p = 0.0156** | -1.2168<br>p = 0.2261 |
| Late fetus/Neonate – Juvenile | -0.7848<br>p = 0.4342 | -0.9381<br>p = 0.3502 | **-2.7371**<br>**p = 0.0074** | -0.4356<br>p = 0.6640 | 1.2254<br>p = 0.2230 | 0.2342<br>p = 0.8153 |
| Juvenile – Adult | -0.4382<br>p = 0.6620 | 0.9021<br>p = 0.3693 | -0.53711<br>p = 0.5922 | **-3.2394**<br>**p = 0.0016** | 1.4342<br>p = 0.1544 | -0.5433<br>p = 0.5879 |

**Table S2.** Pairwise comparison of phenotypic trajectory analysis of skull shape development among genera. Trajectory analysis was performed using the 'trajectory.analysis' [35] function in 'geomorph' [26]. Length, angle and shape of trajectories summarized, including total length of path for each genus. P-values calculated with 1000 permutations. Significant p-values (p<0.05) are in bold. This information is summarized as a heatmap in figure S3.

| | | | | | |
|---|---|---|---|---|---|
| *Absolute difference between path lengths* | | | | | |
| Observed path distances (lengths) by group | | | | | |
| | *Delphinapterus* | *Globicephala* | *Lagenorhynchus* | *Phocoena* | *Stenella* |
| | 0.492 | 0.483 | 0.281 | 0.285 | 0.364 |
| Pairwise absolute differences in path distances, plus statistics | | | | | |
| | d | UCL (95%) | Z | Pr>d | |
| *Delphinapterus:Globicephala* | 0.009 | 0.112 | -1.235 | 0.880 | |
| *Delphinapterus:Lagenorhynchus* | 0.211 | 0.122 | 2.761 | **0.001** | |
| *Delphinapterus:Phocoena* | 0.207 | 0.117 | 2.700 | **0.001** | |
| *Delphinapterus:Stenella* | 0.128 | 0.124 | 1.689 | **0.040** | |
| *Globicephala:Lagenorhynchus* | 0.202 | 0.131 | 2.472 | **0.003** | |
| *Globicephala:Phocoena* | 0.198 | 0.105 | 2.818 | **0.001** | |
| *Globicephala:Stenella* | 0.119 | 0.112 | 1.721 | **0.037** | |
| *Lagenorhynchus:Phocoena* | 0.005 | 0.123 | -1.560 | 0.924 | |
| *Lagenorhynchus:Stenella* | 0.083 | 0.130 | 0.870 | 0.198 | |
| *Phocoena:Stenella* | 0.079 | 0.093 | 1.356 | 0.087 | |
| *Angular differences between trajectory principal axes* | | | | | |
| Pairwise correlations between trajectories, plus statistics | | | | | |
| | r | angle | UCL (95%) | Z | Pr>angle |
| *Delphinapterus:Globicephala* | 0.891 | 26.953 | 22.505 | 2.301 | **0.009** |
| *Delphinapterus:Lagenorhynchus* | 0.751 | 41.328 | 23.860 | 3.777 | **0.001** |
| *Delphinapterus:Phocoena* | 0.901 | 25.667 | 20.261 | 2.612 | **0.006** |
| *Delphinapterus:Stenella* | 0.908 | 24.740 | 20.521 | 2.366 | **0.007** |
| *Globicephala:Lagenorhynchus* | 0.847 | 32.106 | 23.505 | 2.773 | **0.002** |
| *Globicephala:Phocoena* | 0.824 | 34.545 | 20.348 | 3.517 | **0.001** |
| *Globicephala:Stenella* | 0.821 | 34.798 | 20.926 | 3.506 | **0.001** |
| *Lagenorhynchus:Phocoena* | 0.764 | 40.203 | 22.152 | 3.815 | **0.001** |
| *Lagenorhynchus:Stenella* | 0.676 | 47.474 | 22.549 | 4.256 | **0.001** |
| *Phocoena:Stenella* | 0.850 | 31.805 | 18.416 | 3.635 | **0.001** |
| *Shape differences between trajectory vectors* | | | | | |
| Pairwise trajectory shape differences, plus statistics | | | | | |
| | d | UCL (95%) | Z | Pr>d | |
| *Delphinapterus:Globicephala* | 0.315 | 0.502 | 0.374 | 0.359 | |
| *Delphinapterus:Lagenorhynchus* | 0.471 | 0.516 | 1.381 | 0.089 | |
| *Delphinapterus:Phocoena* | 0.270 | 0.469 | 0.090 | 0.479 | |
| *Delphinapterus:Stenella* | 0.355 | 0.466 | 0.851 | 0.213 | |
| *Globicephala:Lagenorhynchus* | 0.178 | 0.500 | -0.975 | 0.828 | |
| *Globicephala:Phocoena* | 0.213 | 0.441 | -0.309 | 0.616 | |
| *Globicephala:Stenella* | 0.181 | 0.447 | -0.735 | 0.758 | |
| *Lagenorhynchus:Phocoena* | 0.277 | 0.454 | 0.257 | 0.394 | |
| *Lagenorhynchus:Stenella* | 0.215 | 0.479 | -0.429 | 0.658 | |
| *Phocoena:Stenella* | 0.188 | 0.398 | -0.400 | 0.654 | |

**Table S3.** Multiple breakpoint test for allometry regression. Model testing was performed using the regression scores obtained from the allometry regression of shape and size (log-transformed Centroid Size – logCS) with genera as a covariate in the package ‘mcp’ [32]. Model formulas and AIC scores of each model and the summary of parameters for the best model are reported. Best model in the first row. Models with an ELPD (log density) difference <4 are considered equally likely. Breakpoints are visualized on the regression for the best model and one breakpoint model in figure S12*a*-*b*.

| **Models’ comparison** | | | |
|---|---|---|---|
| | Model formulas | ELPD_diff | SE_diff |
| One breakpoint model by genus – one breakpoint varying by genus, slope and intercept different among genera | RegScores ~ 1 + (1\|genus) + logCS, RegScores ~ 1 + (1\|genus) ~ 1 + logCS | **0.0** | **0.0** |
| One breakpoint model – one common breakpoint, common slope | RegScores ~ 1 + logCS, ~ 1 + logCS | -30.7 | 5.3 |
| Null model – no breakpoints, common slope | RegScores ~ logCS | -32.8 | 4.7 |
| Null model by genus – no breakpoints, slope and intercept different among genera | RegScores ~ 1 + (1\|genus) + logCS | -32.9 | 4.7 |

**Best model summary**

One breakpoint model by genus

Segments:
1: RegScores ~ 1 + (1 | genus) + logCS
2: RegScores ~ 1 + (1 | genus) ~ 1 + logCS

Family: gaussian (link = 'identity')
Iterations: 9000 from 3 chains

Population-level parameters:

| name | mean | lower | upper | Rhat | n.eff |
|---|---|---|---|---|---|
| cp_1 | 2.505 | 2.400 | 2.642 | 1.1 | 49 |
| cp_1_sd | 1.751 | 0.081 | 4.586 | 1.3 | 372 |
| int_1 | -0.307 | -0.714 | 0.072 | 1.3 | 24 |
| int_2 | -0.211 | -0.260 | -0.154 | 1.1 | 52 |
| logCS_1 | 0.018 | -0.134 | 0.191 | 1.3 | 23 |
| logCS_2 | 0.341 | 0.312 | 0.367 | 1.0 | 418 |
| sigma_1 | 0.027 | 0.021 | 0.032 | 1.0 | 3495 |

**Table S4.** Pairwise comparison of allometry slopes among genera. Allometry models' construction and comparison were calculated using the 'procD.lm' and 'anova' functions in 'geomorph' [26], pairwise comparison was performed using the parse function in the 'RRPP' package [36] on the best model. ANOVA tables of the four tested models (common allometry, different slopes among genera, different intercept among genera divided by pre- and post-natal stages, and different slopes among genera divided by pre- and post-natal stages) are reported, along with model comparison. The last model with different slopes among genera and stages was preferred, and the pairwise analyses of absolute slope distance, angle between slopes and difference in vector lengths for each genus divided in pre- and post-natal stages was conducted. Size is the log-transformed Centroid Size. P-values for pairwise differences calculated with 1000 permutations. Significant p-values (p<0.05) are in bold. Pairwise analysis results are summarized as a heatmap in figure S4*a*-*b*.

| **ANOVA tables tested models** | | | | | | | |
|---|---|---|---|---|---|---|---|
| *Common allometry* | | | | | | | |
| | Df | SS | MS | $R^2$ | F | Z | Pr(>F) |
| size | 1 | 0.757 | 0.757 | 0.468 | 49.294 | 4.350 | **0.001** |
| Residuals | 56 | 0.859 | 0.015 | 0.532 | | | |
| Total | 57 | 1.616 | | | | | |
| *Different slopes genera* | | | | | | | |
| | Df | SS | MS | $R^2$ | F | Z | Pr(>F) |
| size | 1 | 0.757 | 0.757 | 0.468 | 124.175 | 4.973 | **0.001** |
| genus | 4 | 0.477 | 0.119 | 0.295 | 19.585 | 7.262 | **0.001** |
| size:genus | 4 | 0.090 | 0.022 | 0.056 | 3.683 | 5.813 | **0.001** |
| Residuals | 48 | 0.292 | 0.006 | 0.181 | | | |
| Total | 57 | 1.616 | | | | | |
| *Different intercepts genera by stage* | | | | | | | |
| | Df | SS | MS | $R^2$ | F | Z | Pr(>F) |
| size | 1 | 0.757 | 0.757 | 0.468 | 126.690 | 4.959 | **0.001** |
| genus by stage | 9 | 0.579 | 0.064 | 0.358 | 10.770 | 6.886 | **0.001** |
| Residuals | 47 | 0.281 | 0.006 | 0.174 | | | |
| Total | 57 | 1.616 | | | | | |

*Different slopes genera by stage*

| | Df | SS | MS | $R^2$ | F | Z | Pr(>F) |
|---|---|---|---|---|---|---|---|
| size | 1 | 0.757 | 0.757 | 0.468 | 153.641 | 5.059 | **0.001** |
| genus by stage | 9 | 0.579 | 0.064 | 0.358 | 13.061 | 7.204 | **0.001** |
| size:genus by stage | 9 | 0.094 | 0.010 | 0.058 | 2.111 | 5.715 | **0.001** |
| Residuals | 38 | 0.187 | 0.005 | 0.116 | | | |
| Total | 57 | 1.616 | | | | | |

**ANOVA models' comparison**

| | ResDf | Df | RSS | SS | MS | $R^2$ | F | Z | P |
|---|---|---|---|---|---|---|---|---|---|
| Common allometry (Null) | 56 | 1 | 0.859 | 0.000 | | | | | |
| Different slopes genera | 48 | 8 | 0.292 | 0.567 | 0.071 | 0.351 | 11.634 | 7.746 | **0.001** |
| Different intercepts genera by stage | 47 | 9 | 0.281 | 0.579 | 0.064 | 0.358 | 10.770 | 6.886 | **0.001** |
| Different slopes genera by stage | 38 | 18 | 0.187 | 0.672 | 0.037 | 0.416 | 7.586 | 9.272 | **0.001** |
| Total | 57 | 1.616 | | | | | | | |

**Pairwise comparisons genera by stage**

| | *Absolute distances between slopes* | | | *Angular differences between slopes* | | | *Differences in slope vector length* | | |
|---|---|---|---|---|---|---|---|---|---|
| | d | Z | Pr>d | angle | Z | Pr>angle | d | Z | Pr>d |
| *Delphinapterus*_postnatal:*Delphinapterus*_ prenatal | 1.018 | -0.947 | 0.829 | 77.748 | 0.378 | 0.365 | 0.645 | -1.343 | 0.908 |
| *Delphinapterus*_postnatal:*Globicephala*_ postnatal | 1.070 | -0.857 | 0.804 | 76.014 | -0.089 | 0.539 | 0.392 | -2.019 | 0.984 |
| *Delphinapterus*_postnatal:*Globicephala*_ prenatal | 0.998 | -1.062 | 0.859 | 75.609 | 0.222 | 0.413 | 0.711 | -0.960 | 0.836 |
| *Delphinapterus*_postnatal:*Lagenorhynchus* _postnatal | 1.406 | 0.033 | 0.477 | 81.390 | -0.099 | 0.540 | 0.097 | -1.670 | 0.946 |
| *Delphinapterus*_postnatal:*Lagenorhynchus* _prenatal | 1.037 | -0.993 | 0.839 | 79.076 | 0.207 | 0.416 | 0.595 | -1.091 | 0.857 |
| *Delphinapterus*_postnatal:*Phocoena*_ postnatal | 1.151 | -2.374 | 0.994 | 69.672 | -1.257 | 0.897 | 0.042 | -1.327 | 0.893 |
| *Delphinapterus*_postnatal:*Phocoena*_ prenatal | 1.028 | -0.917 | 0.824 | 79.174 | 0.428 | 0.352 | 0.642 | -1.247 | 0.887 |
| *Delphinapterus*_postnatal:*Stenella*_ postnatal | 1.063 | -1.071 | 0.855 | 65.745 | -1.166 | 0.868 | 0.105 | -3.396 | 1.000 |

| | | | | | | | | | |
|---|---|---|---|---|---|---|---|---|---|
| *Delphinapterus*_postnatal:*Stenella*_prenatal | 1.006 | -1.025 | 0.842 | 75.818 | 0.208 | 0.415 | 0.637 | -1.347 | 0.910 |
| *Delphinapterus*_prenatal:*Globicephala*_postnatal | 0.685 | 3.370 | **0.001** | 80.269 | 2.833 | **0.002** | 0.253 | 1.633 | **0.049** |
| *Delphinapterus*_prenatal:*Globicephala*_prenatal | 0.148 | 2.167 | **0.017** | 21.846 | 1.792 | **0.035** | 0.066 | 2.027 | **0.015** |
| *Delphinapterus*_prenatal:*Lagenorhynchus*_postnatal | 1.036 | 1.907 | **0.033** | 66.745 | 0.293 | 0.388 | 0.742 | 1.747 | **0.036** |
| *Delphinapterus*_prenatal:*Lagenorhynchus*_prenatal | 0.302 | 0.539 | 0.302 | 42.967 | 0.588 | 0.275 | 0.050 | -0.405 | 0.663 |
| *Delphinapterus*_prenatal:*Phocoena*_postnatal | 0.994 | -0.891 | 0.794 | 80.035 | 0.492 | 0.330 | 0.603 | -1.310 | 0.912 |
| *Delphinapterus*_prenatal:*Phocoena*_prenatal | 0.211 | 2.419 | **0.008** | 31.851 | 2.129 | **0.013** | 0.003 | -1.540 | 0.931 |
| *Delphinapterus*_prenatal:*Stenella*_postnatal | 0.807 | 3.466 | **0.001** | 60.571 | 1.000 | 0.164 | 0.540 | 3.106 | **0.001** |
| *Delphinapterus*_prenatal:*Stenella*_prenatal | 0.203 | 2.751 | **0.002** | 30.329 | 2.505 | **0.009** | 0.008 | -0.839 | 0.780 |
| *Globicephala*_postnatal:*Globicephala*_prenatal | 0.671 | 3.287 | **0.001** | 82.092 | 2.910 | **0.002** | 0.320 | 2.230 | **0.011** |
| *Globicephala*_postnatal:*Lagenorhynchus*_postnatal | 0.921 | 0.927 | 0.185 | 54.957 | -1.417 | 0.922 | 0.489 | 0.977 | 0.156 |
| *Globicephala*_postnatal:*Lagenorhynchus*_prenatal | 0.783 | 3.247 | **0.001** | 92.107 | 2.823 | **0.001** | 0.204 | 1.582 | 0.050 |
| *Globicephala*_postnatal:*Phocoena*_postnatal | 1.102 | -0.534 | 0.697 | 82.527 | 0.330 | 0.365 | 0.349 | -1.958 | 0.973 |
| *Globicephala*_postnatal:*Phocoena*_prenatal | 0.751 | 3.481 | **0.001** | 91.138 | 3.176 | **0.001** | 0.250 | 1.658 | **0.041** |
| *Globicephala*_postnatal:*Stenella*_postnatal | 0.780 | 2.309 | **0.008** | 56.554 | -0.509 | 0.688 | 0.286 | 1.842 | **0.022** |
| *Globicephala*_postnatal:*Stenella*_prenatal | 0.722 | 3.482 | **0.001** | 85.835 | 3.054 | **0.001** | 0.245 | 1.555 | 0.053 |
| *Globicephala*_prenatal:*Lagenorhynchus*_postnatal | 1.064 | 2.026 | **0.025** | 70.856 | 0.593 | 0.281 | 0.808 | 2.044 | **0.020** |
| *Globicephala*_prenatal:*Lagenorhynchus*_prenatal | 0.295 | 0.390 | 0.353 | 42.931 | 0.554 | 0.297 | 0.116 | 0.670 | 0.279 |
| *Globicephala*_prenatal:*Phocoena*_postnatal | 0.989 | -0.927 | 0.814 | 81.337 | 0.559 | 0.293 | 0.669 | -0.933 | 0.838 |
| *Globicephala*_prenatal:*Phocoena*_prenatal | 0.214 | 2.341 | **0.008** | 33.619 | 2.288 | **0.014** | 0.070 | 1.385 | 0.082 |
| *Globicephala*_prenatal:*Stenella*_postnatal | 0.839 | 3.653 | **0.001** | 64.900 | 1.298 | 0.106 | 0.606 | 3.539 | **0.001** |
| *Globicephala*_prenatal:*Stenella*_prenatal | 0.226 | 3.248 | **0.001** | 35.236 | 3.148 | **0.001** | 0.074 | 1.900 | **0.026** |

| | | | | | | | | | |
|---|---|---|---|---|---|---|---|---|---|
| *Lagenorhynchus*_postnatal: *Lagenorhynchus*_prenatal | 1.162 | 2.206 | **0.017** | 83.888 | 1.141 | 0.123 | 0.693 | 1.856 | **0.028** |
| *Lagenorhynchus*_postnatal:*Phocoena*_ postnatal | 1.305 | -0.242 | 0.596 | 76.053 | -0.498 | 0.703 | 0.140 | -1.281 | 0.894 |
| *Lagenorhynchus*_postnatal:*Phocoena*_ prenatal | 1.105 | 2.142 | **0.018** | 77.046 | 0.940 | 0.189 | 0.739 | 1.777 | **0.034** |
| *Lagenorhynchus*_postnatal:*Stenella*_ postnatal | 0.767 | -0.272 | 0.596 | 42.586 | -2.777 | 0.998 | 0.203 | -0.255 | 0.617 |
| *Lagenorhynchus*_postnatal:*Stenella*_ prenatal | 1.058 | 1.987 | **0.024** | 70.044 | 0.496 | 0.317 | 0.734 | 1.741 | **0.040** |
| *Lagenorhynchus*_prenatal:*Phocoena*_ postnatal | 1.064 | -0.661 | 0.730 | 88.162 | 0.816 | 0.216 | 0.553 | -1.070 | 0.865 |
| *Lagenorhynchus*_prenatal:*Phocoena*_ prenatal | 0.286 | -0.115 | 0.547 | 40.373 | -0.104 | 0.538 | 0.046 | -0.289 | 0.607 |
| *Lagenorhynchus*_prenatal:*Stenella*_ postnatal | 0.947 | 3.978 | **0.001** | 79.766 | 1.791 | **0.035** | 0.490 | 2.666 | **0.001** |
| *Lagenorhynchus*_prenatal:*Stenella*_ prenatal | 0.310 | 0.513 | 0.307 | 43.823 | 0.522 | 0.306 | 0.042 | -0.521 | 0.694 |
| *Phocoena*_postnatal:*Phocoena*_prenatal | 1.001 | -0.879 | 0.796 | 80.987 | 0.507 | 0.310 | 0.599 | -1.229 | 0.900 |
| *Phocoena*_postnatal:*Stenella*_postnatal | 1.028 | -1.067 | 0.851 | 65.119 | -1.136 | 0.870 | 0.063 | -3.629 | 1.000 |
| *Phocoena*_postnatal:*Stenella*_prenatal | 0.977 | -1.003 | 0.833 | 77.212 | 0.280 | 0.394 | 0.595 | -1.335 | 0.916 |
| *Phocoena*_prenatal:*Stenella*_postnatal | 0.845 | 3.820 | **0.001** | 66.245 | 1.280 | 0.117 | 0.536 | 3.040 | **0.001** |
| *Phocoena*_prenatal:*Stenella*_prenatal | 0.186 | 0.957 | 0.160 | 27.731 | 0.672 | 0.250 | 0.005 | -1.481 | 0.920 |
| *Stenella*_postnatal:*Stenella*_prenatal | 0.790 | 3.385 | **0.001** | 58.197 | 0.715 | 0.236 | 0.532 | 2.967 | **0.002** |

**Table S5.** Pairwise comparison of allometry slopes ancestral states and genera. A regression model was calculated with the ‘lm’ function in base R [21] recreating the best model found for extant taxa in the allometry analysis (formula = RegScores ~ logCS * genus/node by stage). The regression scores and logCS for extant taxa were extracted from the allometry model of size with slopes varying among genera by stage. Regression scores for the ancestral nodes was calculated based on the ancestral intercepts and slopes estimate with the ‘fastAnc’ function in the package ‘phytools’ [37] using a distribution of logCS values based on the range of modern taxa. Each node divided in pre- and post-natal was considered as a group to be compared with the extant genera. Nodes are numbered as follows: 1 – Delphinoidea, 2 – Monodontidae+Phocoenidae, 3 – Delphinidae, 4 – *Stenella*+*Globicephala*. Pairwise comparison was performed in the package ‘emmeans’ [34]. Upper triangle: Tukey-adjusted P-values of mean comparisons, Diagonal: emmean estimates for each group (italicized), Lower triangle: comparisons of emmean estimates earlier vs. later. Pairwise comparisons among extant genera are shaded as the values obtained using the pairwise function on the original procD.lm model are more reliable tant these emmeans values, and they are not illustrated in the heatmap plots (figure S6*a*-*b*). Significant p-values (p<0.05) are in bold.

| | 1_ post | 1_ pre | 2_ post | 2_ pre | 3_ post | 3_ pre | 4_ post | 4_ pre | Delp_ post | Delp_ pre | Glob_ post | Glob_ pre | Lage_ post | Lage_ pre | Phoc_ post | Phoc_ pre | Sten_ post | Sten_ pre |
|---|---|---|---|---|---|---|---|---|---|---|---|---|---|---|---|---|---|---|
| 1_ post | *0.016* | 0.670 | 1.000 | **0.006** | 0.993 | 1.000 | 1.000 | 1.000 | 1.000 | **<0.0001** | 1.000 | **<0.0001** | 0.764 | **0.007** | 0.920 | 0.950 | **0.001** | **<0.0001** |
| 1_ pre | 0.011 | *0.006* | 0.070 | 0.557 | 1.000 | **0.016** | 0.078 | 0.068 | 1.000 | **<0.0001** | 1.000 | **<0.0001** | 0.960 | **<0.0001** | 0.125 | 1.000 | **<0.0001** | **<0.0001** |
| 2_ post | -0.005 | -0.016 | *0.022* | **<0.0001** | 0.625 | 1.000 | 1.000 | 1.000 | 1.000 | **<0.0001** | 1.000 | **<0.0001** | 0.602 | 0.053 | 0.998 | 0.554 | **0.010** | **<0.0001** |
| 2_ pre | 0.019 | 0.008 | 0.024 | *-0.003* | 0.609 | **<0.0001** | **<0.0001** | **<0.0001** | 1.000 | **<0.0001** | 1.000 | **<0.0001** | 0.997 | **<0.0001** | **0.005** | 1.000 | **<0.0001** | **<0.0001** |
| 3_ post | 0.008 | -0.003 | 0.013 | -0.011 | *0.008* | 0.661 | 0.648 | 0.871 | 1.000 | **<0.0001** | 1.000 | **<0.0001** | 0.935 | **<0.0001** | 0.358 | 1.000 | **<0.0001** | **<0.0001** |
| 3_ pre | -0.003 | -0.013 | 0.002 | -0.022 | -0.011 | *0.019* | 1.000 | 1.000 | 1.000 | **<0.0001** | 1.000 | **<0.0001** | 0.667 | **0.008** | 0.969 | 0.639 | **0.001** | **<0.0001** |
| 4_ post | -0.005 | -0.016 | 0.000 | -0.024 | -0.013 | -0.002 | *0.021* | 1.000 | 1.000 | **<0.0001** | 1.000 | **<0.0001** | 0.608 | 0.050 | 0.997 | 0.572 | **0.009** | **<0.0001** |
| 4_ pre | -0.001 | -0.012 | 0.004 | -0.020 | -0.009 | 0.002 | 0.004 | *0.018* | 1.000 | **<0.0001** | 1.000 | **<0.0001** | 0.717 | **0.004** | 0.924 | 0.802 | **0.001** | **<0.0001** |
| Delp_ post | 0.001 | -0.010 | 0.006 | -0.018 | -0.007 | 0.003 | 0.006 | 0.002 | *0.016* | 1.000 | 1.000 | 1.000 | 1.000 | 1.000 | 1.000 | 1.000 | 1.000 | 1.000 |
| Delp_ pre | 0.050 | 0.040 | 0.055 | 0.031 | 0.042 | 0.053 | 0.055 | 0.051 | 0.049 | *-0.034* | 0.425 | 1.000 | 1.000 | **<0.0001** | **<0.0001** | **<0.0001** | **<0.0001** | **<0.0001** |
| Glob_ post | 0.002 | -0.008 | 0.007 | -0.017 | -0.006 | 0.005 | 0.007 | 0.003 | 0.002 | -0.048 | *0.014* | 0.407 | 0.973 | 0.882 | 1.000 | 1.000 | 0.701 | 0.252 |
| Glob_ pre | 0.050 | 0.040 | 0.056 | 0.031 | 0.042 | 0.053 | 0.055 | 0.052 | 0.050 | 0.000 | 0.048 | *-0.034* | 1.000 | **<0.0001** | **<0.0001** | **<0.0001** | **<0.0001** | **<0.0001** |
| Lage_ post | 0.049 | 0.038 | 0.054 | 0.030 | 0.041 | 0.051 | 0.054 | 0.050 | 0.048 | -0.002 | 0.046 | -0.002 | *-0.032* | **0.040** | 0.323 | 0.982 | **0.018** | **0.002** |
| Lage_ pre | -0.035 | -0.046 | -0.030 | -0.054 | -0.043 | -0.033 | -0.030 | -0.034 | -0.036 | -0.085 | -0.038 | -0.086 | -0.084 | *0.052* | 0.978 | **<0.0001** | 1.000 | 0.959 |
| Phoc_ post | -0.017 | -0.028 | -0.012 | -0.036 | -0.025 | -0.014 | -0.012 | -0.016 | -0.018 | -0.067 | -0.019 | -0.068 | -0.066 | 0.018 | *0.034* | 0.260 | 0.801 | **0.045** |
| Phoc_ pre | 0.013 | 0.002 | 0.018 | -0.006 | 0.005 | 0.015 | 0.018 | 0.014 | 0.012 | -0.038 | 0.010 | -0.038 | -0.036 | 0.048 | 0.030 | *0.004* | **<0.0001** | **<0.0001** |
| Sten_ post | -0.042 | -0.052 | -0.037 | -0.061 | -0.050 | -0.039 | -0.037 | -0.041 | -0.042 | -0.092 | -0.044 | -0.092 | -0.090 | -0.006 | -0.025 | -0.054 | *0.058* | 1.000 |
| Sten_ pre | -0.051 | -0.061 | -0.045 | -0.070 | -0.059 | -0.048 | -0.046 | -0.049 | -0.051 | -0.101 | -0.053 | -0.101 | -0.099 | -0.015 | -0.033 | -0.063 | -0.009 | *0.067* |

**Table S6.** ANOVA analyses of the first two PC components for raw shape data and allometry residuals. PC components obtained from the ‘gm.prcomp’ function in ‘geomorph’ [26], ‘lm’ function in base R [21] used to conduct ANOVA analyses. Four independent variables were tested for raw data: size (log-transformed centroid size), genera, growth category and asymmetry (distance from symmetric shape). Only genera and asymmetry were tested for residuals as the allometry correction eliminates the effect of size and growth stage from the data. Asymmetry and size results reported as ANOVA table (continuous variables), growth category and genera as summary table (categorical variables). Significant p-values (p<0.05) are in bold.

| **Raw data** | | | | | | | | | |
|---|---|---|---|---|---|---|---|---|---|
| PC1 | | | | | PC2 | | | | |
| *ANOVA table size* | | | | | *ANOVA table size* | | | | |
| | Df | Sum_Sq | Mean_Sq | F | Pr(>F) | Df | Sum_Sq | Mean_Sq | F | Pr(>F) |
| *size* | 1 | 0.71474 | 0.71474 | 194.71 | **< 2.2e-16** | *size* 1 | 0.039659 | 0.039659 | 11.384 | **0.00135** |
| Residuals | 56 | 0.20556 | 0.00367 | | | Residuals 56 | 0.195086 | 0.003484 | | |
| *ANOVA table asymmetry* | | | | | *ANOVA table asymmetry* | | | | |
| | Df | Sum_Sq | Mean_Sq | F | Pr(>F) | Df | Sum_Sq | Mean_Sq | F | Pr(>F) |
| *asymmetry* | 1 | 0.26039 | 0.260390 | 22.096 | **1.731e-05** | *asymmetry* 1 | 0.058508 | 0.058508 | 18.591 | **6.628e-05** |
| Residuals | 56 | 0.65992 | 0.011784 | | | Residuals 56 | 0.176237 | 0.003147 | | |
| *Summary table genera* | | | | | *Summary table genera* | | | | |
| Genera | Estimate | Std. Error | t value | Pr(>\|t\|) | Genera | Estimate | Std. Error | t value | Pr(>\|t\|) |
| *Delphinapterus* (ref.) | -0.02987 | 0.04040 | -0.739 | 0.4631 | *Delphinapterus* (ref.) | -0.08839 | 0.01090 | -8.110 | **7.48e-11** |
| *Globicephala* | -0.01181 | 0.05569 | -0.212 | 0.8329 | *Globicephala* | 0.02521 | 0.01502 | 1.678 | 0.0993 |
| *Lagenorhynchus* | 0.07990 | 0.05890 | 1.357 | 0.1807 | *Lagenorhynchus* | 0.10596 | 0.01589 | 6.669 | **1.54e-08** |
| *Phocoena* | -0.01767 | 0.05111 | -0.346 | 0.7309 | *Phocoena* | 0.10959 | 0.01379 | 7.949 | **1.35e-10** |
| *Stenella* | 0.09226 | 0.05051 | 1.827 | 0.0734 | *Stenella* | 0.14892 | 0.01362 | 10.931 | **3.41e-15** |

*Summary table growth <u>categories</u>* (left)

| Growth categories | Estimate | Std. Error | t value | Pr(>\|t\|) |
|---|---|---|---|---|
| Early fetus (ref.) | -0.15737 | 0.01630 | -9.654 | **2.34e-13** |
| Late fetus/neonate | 0.16799 | 0.02341 | 7.176 | **2.14e-09** |
| Juvenile | 0.24878 | 0.02426 | 10.256 | **2.77e-14** |
| Adult | 0.26879 | 0.02601 | 10.335 | **2.10e-14** |

*Summary table growth <u>categories</u>* (right)

| Growth categories | Estimate | Std. Error | t value | Pr(>\|t\|) |
|---|---|---|---|---|
| Early fetus (ref.) | 0.013351 | 0.015358 | 0.869 | 0.3885 |
| Late fetus/neonate | -0.001649 | 0.022056 | -0.075 | 0.9407 |
| Juvenile | -0.016031 | 0.022853 | -0.701 | 0.4860 |
| Adult | -0.047592 | 0.024503 | -1.942 | 0.0573 |

**Allometry residuals**

<u>PC1</u>

*ANOVA table <u>asymmetry</u>*

| | Df | Sum_Sq | Mean_Sq | F | Pr(>F) |
|---|---|---|---|---|---|
| *asymmetry* | 1 | 0.00656 | 0.006557 | 1.0093 | 0.3194 |
| Residuals | 56 | 0.36384 | 0.006497 | | |

*Summary table <u>genera</u>*

| Genera | Estimate | Std. Error | t value | Pr(>\|t\|) |
|---|---|---|---|---|
| *Delphinapterus* (ref.) | -0.095800 | 0.011100 | -8.630 | **1.12e-11** |
| *Globicephala* | 0.006884 | 0.015301 | 0.450 | 0.655 |
| *Lagenorhynchus* | 0.128603 | 0.016182 | 7.948 | **1.36e-10** |
| *Phocoena* | 0.094810 | 0.014041 | 6.752 | **1.13e-08** |
| *Stenella* | 0.189788 | 0.013876 | 13.678 | **< 2e-16** |

<u>PC2</u>

*ANOVA table <u>asymmetry</u>*

| | Df | Sum_Sq | Mean_Sq | F | Pr(>F) |
|---|---|---|---|---|---|
| *asymmetry* | 1 | 0.037353 | 0.037353 | 28.476 | **1.775e-06** |
| Residuals | 56 | 0.073459 | 0.001312 | | |

*Summary table <u>genera</u>*

| Genera | Estimate | Std. Error | t value | Pr(>\|t\|) |
|---|---|---|---|---|
| *Delphinapterus* (ref.) | -0.021502 | 0.008685 | -2.476 | **0.0165** |
| *Globicephala* | 0.0006087 | 0.011972 | 0.051 | 0.9596 |
| *Lagenorhynchus* | 0.0122680 | 0.012661 | 0.969 | 0.3370 |
| *Phocoena* | 0.0816567 | 0.010986 | 7.433 | **9.08e-10** |
| *Stenella* | -0.005123 | 0.010857 | -0.47 | 0.6390 |

**Table S7.** Landmarks (LM), curves and number of semilandmarks (sLMs) used in this study. Left landmarks and curves are italicized, points on the midline are in bold. Landmarks and curves are pictured with fixed landmarks numbered in figure S8.

| Bone | LM | Landmark definition | Curve | Curve definition | No. sLMs |
|---|---|---|---|---|---|
| maxilla | *1*, 10 | anterior end maxilla | *1-2*, 10-11 | lateral edge of rostral portion of maxilla | 25 |
| maxilla | *2*, 11 | antorbital notch - deepest point | *2-3*, 11-12 | anterior margin antorbital notch | 5 |
| maxilla | *3*, 12 | lateral edge of antorbital notch (maxilla) | *3-4*, 12-13 | lateral and posterior edge ascending process maxilla | 20 |
| maxilla | *4*, 13 | medial end ascending process of maxilla (dorsalmost point of contact with nasal/ethmoid) | — | — | — |
| maxilla | *5*, 14 | posterior end premaxilla (contact on maxilla) | *1-5*, 10-14 | lateral edge premaxilla on maxilla | 25 |
| nasal | *6*, 15 | ventral/anterior medial end nasal | *6-8*, 15-17 | ventrolateral edge of nasal | 5 |
| nasal | *7*, 16 | dorsal/posterior medial end nasal | *8-6*, 17-15 | dorsomedial edge of nasal | 5 |
| nasal | *8*, 17 | dorsal/posterior lateral end nasal | — | — | — |
| nasal | *9*, 18 | ventral/anterior lateral end nasal | — | — | — |
| premaxilla | *19*, 20 | anterior end premaxilla | — | — | — |
| frontal | *21*, 28 | ventral end preorbital process of frontal | *21-22*, 28-29 | orbital process of frontal | 7 |
| frontal | *22*, 29 | ventral end postorbital process of frontal | — | — | — |
| squamosal | *23*, 30 | anterior end supramastoid crest of squamosal | *23-24*, 30-31 | anterior edge of squamosal | 3 |
| squamosal | *24*, 31 | anterior end zygomatic process of squamosal | *24-25*, 31-32 | ventral edge of squamosal | 5 |
| squamosal | *25*, 32 | ventral end postglenoid process of squamosal | *25-26*, 32-33 | posterior edge of squamosal | 10 |
| squamosal | *26*, 33 | posterior end supramastoid crest of squamosal | *26-23*, 33-30 | dorsal edge of squamosal | 5 |
| exoccipital | *27*, 34 | lateralmost edge of exoccipital (contact -present or future- with posteriormost edge of squamosal plate) | *35-27*, 39-34 | lateral edge of exoccipital | 7 |
| exoccipital | *35*, 39 | ventral end paraoccipital process of exoccipital | — | — | — |
| occipital condyle | *36*, 40 | dorsal end occipital condyle | *37-36*, 41-40 | lateral edge of occipital condyle | 7 |
| occipital condyle | *37*, 41 | ventral end occipital condyle | *36-37*, 40-41 | medial edge of occipital condyle | 7 |
| basioccipital | *38*, 42 | posterior end of the basioccipital crest | *53-38*, 53-42 | posterior edge of basioccipital | 5 |
| maxilla | *48*, 43 | anterior end tooth row on maxilla | *48-49*, 43-44 | medial edge of tooth row on maxilla | 20 |
| maxilla | *49*, 44 | posterior end tooth row on maxilla | — | — | — |
| jugal | *50*, 45 | anterior end maxillary process of jugal (anteriormost point of contact between jugal and lacrimal/frontal) | — | — | — |
| palatine | *51*, 46 | anteriormost point of palatine crest, edge of greater palatal foramen | *51-52*, 46-47 | medial edge of palatine | 9 |
| palatine | *52*, 47 | posteriormost point of nasal spine of palatine | — | — | — |
| basioccipital | *56*, 54 | anteriormost lateral edge of basioccipital | — | — | — |

| | | | | | |
|---|---|---|---|---|---|
| interparietal | *57*, 59 | anterolateral edge of interparietal (contact -present or future- with frontal) | — | — | — |
| supraoccipital | *60*, 62 | lateral edge of supraoccipital (contact -present or future- with interparietal or limit of medial part of suproccipital if interparietal obscured) | — | — | — |
| basioccipital | **53** | posteriormost edge of basioccipital | *38-56*, 42-54 | lateral edge of basioccipital | 10 |
| basioccipital | **55** | anteriormost edge of basioccipital | *56-55*, 54-55 | anterior edge of basioccipital | 5 |
| interparietal | **58** | midpoint anterior margin of interparietal (contact with frontal) | *57-58*, 59-58 | anterior edge of interparietal (contact with frontal) | 6 |
| supraoccipital | **61** | midpoint anterior margin of supraoccipital (contact with interparietal), dorsal end of supraoccipital crest | *60-61*, 62-61 | dorsal edge of supraoccipital (contact with interparietal) | 5 |

**Table S8.** Pairwise comparison of mean distances between DA and symmetric shape among consecutive growth stages for entire dataset and by taxon, all fixed landmarks included. Distance between asymmetric and symmetric shape for each specimen calculated using 'landVR' package [29] and then analysed grouped by growth stage. Difference between means (t) and p-value obtained with a Welch Two Sample t-test ('t.test' function) are reported for each pair. Significant p-values ($p<0.05$) are in bold.

| | Entire dataset | *Delphinapterus* | *Globicephala* | *Lagenorhynchus* | *Phocoena* | *Stenella* |
|---|---|---|---|---|---|---|
| Early fetus – Late fetus/neonate | **-2.9599**<br>**p = 0.0037** | **-7.0063**<br>**p = 3.55e-10** | 0.1422<br>p = 0.8871 | 0.8542<br>p = 0.3951 | **-2.1958**<br>**p = 0.0299** | -1.1695<br>p = 0.2444 |
| Late fetus/Neonate – Juvenile | -1.6271<br>p = 0.1069 | -1.7231<br>p = 0.0887 | **-3.1195**<br>**p = 0.0024** | -0.4344<br>p = 0.6648 | 1.4148<br>p = 0.1597 | -0.6667<br>p = 0.5063 |
| Juvenile – Adult | -1.2087<br>p = 0.23 | 0.7976<br>p = 0.4268 | -1.7016<br>p = 0.0929 | **-3.1948**<br>**p = 0.002** | 1.2025<br>p = 0.2316 | -1.4349<br>p = 0.1552 |

**Dataset S1 (separate file).** List of specimens analysed in the study. This table includes specimens' codes used to identify specimens in analyses, museum accession number, taxonomic information, total length and age estimate, growth category assignment, specimen preparation, scan type, as well as reference to who provided the data. Additional notes are provided in a separate sheet.

**Dataset S2 (separate file).** Details on acquisition specimens digitized using CT scanning. Facility, CT system model used and details on image acquisition (kV, mA, slice thickness, voxel size, number of images, output format for images) are reported, along with other details when available. Additional notes are provided in a separate sheet.

### Supplemental references


[1] Roston, R.A. & Roth, V.L. 2019 Cetacean skull telescoping brings evolution of cranial sutures into focus. *Anat. Rec.* **302**, 1055-1073. (doi:10.1002/ar.24079).
[2] Berta, A. & Lanzetti, A. 2020 Feeding in marine mammals: an integration of evolution and ecology through time. *Palaeontol. Electron.* **23**, a40. (doi:doi.org/10.26879/951).
[3] Vollmer, N.L., Ashe, E., Brownell, R.L., Cipriano, F., Mead, J.G., Reeves, R.R., Soldevilla, M.S. & Williams, R. 2019 Taxonomic revision of the dolphin genus *Lagenorhynchus*. *Mar. Mamm. Sci.* **35**, 957-1057. (doi:10.1111/mms.12573).
[4] McGowen, M.R., Tsagkogeorga, G., Álvarez-Carretero, S., dos Reis, M., Struebig, M., Deaville, R., Jepson, P.D., Jarman, S., Polanowski, A., Morin, P.A., et al. 2020 Phylogenomic resolution of the cetacean tree of life using target sequence capture. *Syst. Biol.* **69**, 479-501. (doi:10.1093/sysbio/syz068).
[5] Pyenson, N.D. & Sponberg, S.N. 2011 Reconstructing body size in extinct crown Cetacea (Neoceti) using allometry, phylogenetic methods and tests from the fossil record. *J. Mamm. Evol.* **18**, 269. (doi:10.1007/s10914-011-9170-1).
[6] Frazer, J.F.D. & Huggett, A.S.G. 1973 Specific foetal growth rates of cetaceans. *J. Zool.* **169**, 111-126. (doi:10.1111/j.1469-7998.1973.tb04656.x).
[7] Laws, R.M. 1959 The foetal growth rates of whales with special reference to the fin whale, *Balaenoptera physalus* Linn. *Disc. Rep.* **29**, 281-308.
[8] Sterba, O., Klima, M. & Schildger, B. 2000 *Embryology of dolphins: staging and ageing of embryos and fetuses of some cetaceans*. New York, NY, Springer-Verlag; 133 p.
[9] Perrin, W.F. & Reilly, S.B. 1984 Reproductive parameters of dolphins and small whales of the family Delphinidae.In *Rep Int Whal Comm*, pp. 97-133, Special Issue 6. La Jolla, CA.
[10] Lanzetti, A., Berta, A. & Ekdale, E.G. 2020 Prenatal development of the humpback whale: growth rate, tooth loss and skull shape changes in an evolutionary framework. *Anat. Rec.* **303**, 180-204. (doi:10.1002/ar.23990).
[11] Bernard, H.J. & Reilly, S.B. 1998 Pilot whales - *Globicephala* Lesson, 1828. In *Handbook of marine mammals: the second book of dolphins and the porpoises* (eds. S.H. Ridgway & R. Harrison), pp. 245-280. London, Academic Press.
[12] Reeves, R.R., Smeenk, C., Brownell Jr, R.L. & Kinze, C.C. 1998 Atlantic White-sided dolphin - *Lagenorhynchus acutus* (Gray, 1828). In *Handbook of marine mammals: the second book of dolphins and the porpoises* (eds. S.H. Ridgway & R. Harrison), pp. 31-56. London, Academic Press.
[13] Read, A.J. 1999 Harbour porpoise - *Phocoena phocoena* (Linnaeus, 1758). In *Handbook of marine mammals: the second book of dolphins and the porpoises* (eds. S.H. Ridgway & R. Harrison), pp. 323-356. London, Academic Press.
[14] Perrin, W.F., Coe, J.M. & Zweifel, J.R. 1976 Growth and reproduction of the spotted porpoise, *Stenella attenuata*, in the offshore eastern tropical pacific. *Fish. Bull.* **74**, 229-269.
[15] Perrin, W.F., Hole, D.B. & Miller, R.B. 1977 Growth and reproduction of the Eastern spinner dolphin, a geographical form of *Stenella longirostris* in the Eastern Tropical Pacific. *Fish. Bull.* **75**, 725-750.

[16] Stewart, B.E. & Stewart, R.E.A. 1989 *Delphinapterus leucas*. *Mamm. Species* **336**, 1-8.
[17] Tomilin, A.G. 1967 *Mammals of the USSR and adjacent countries, Volume IX, Cetacea*. Jerusalem, Israel Program for Scientific Translations; 717 p.
[18] Gower, R.J.B., Vincent, F., Sterling, J.N., João Vasco, L. & David, J. 2022 A new pseudosuchian archosaur, *Mambawakale ruhuhu* gen. et sp. nov., from the Middle Triassic Manda Beds of Tanzania. *Roy. Soc. Open Sci.* **9**, 211622. (doi:10.1098/rsos.211622).
[19] Schneider, C.A., Rasband, W.S. & Eliceiri, K.W. 2012 NIH Image to ImageJ: 25 years of image analysis. *Nat. Methods* **9**, 671-675. (doi:10.1038/nmeth.2089).
[20] Bookstein, F.L. 1991 *Morphometric tools for landmark data: geometry and biology*. Cambridge, Cambridge University Press; 435 p.
[21] R Core Team. 2021 R: A language and environment for statistical computing. See http://www.r-project.org.
[22] Felice, R.N. 2021 SURGE: SURface geometric morphometrics for everyone. See https://github.com/rnfelice/SURGE.
[23] Bardua, C., Felice, R.N., Watanabe, A., Fabre, A.-C. & Goswami, A. 2019 A practical guide to sliding and surface semilandmarks in morphometric analyses. *Integr. Org. Biol.* **1**, obz016. (doi:10.1093/iob/obz016).
[24] Schlager, S. 2013 Morpho: calculations and visualizations related to geometric morphometrics. See http://cran.r-project.org/package=morpho.
[25] Schlager, S. 2017 Morpho and rvcg – shape analysis in R: R-packages for geometric morphometrics, shape analysis and surface manipulations. In *Statistical Shape and Deformation Analysis* (eds. G. Zheng, S. Li & G. Szekely), pp. 217-256. London, Academic Press.
[26] Baken, E.K., Collyer, M.L., Kaliontzopoulou, A. & Adams, D.C. 2021 geomorph v4.0 and gmShiny: enhanced analytics and a new graphical interface for a comprehensive morphometric experience. *Methods Ecol. Evol.* **12**, 2355-2363. (doi:10.1111/2041-210x.13723).
[27] Huggenberger, S., Leidenberger, S. & Oelschläger, H.H.A. 2017 Asymmetry of the nasofacial skull in toothed whales (Odontoceti). *J. Zool.* **302**, 15-23. (doi:10.1111/jzo.12425).
[28] Kuzmin, A.A. 1976 Embryogenesis of the osseus skull of the sperm whale. In *Investigations on Cetacea* (ed. G. Pilleri), pp. 187-202. Berne, Switzerland, Institute of Brain Anatomy, University of Berne.
[29] Guillerme, T., Weisbecker, V. & Marcy, A.E. 2019 landvR: Tools for measuring landmark position variation. See https://doi.org/10.5281/zenodo.2620785.
[30] Mead, J.G. & Fordyce, R.E. 2009 *The therian skull: a lexicon with emphasis on the odontocetes*. Washington, D.C., Smithsonian institution press; 249 p.
[31] Murdoch, D. & Adler, D. 2021 rgl: 3D visualization using OpenGL. See https://CRAN.R-project.org/package=rgl.
[32] Lindeløv, J.K. 2020 mcp: An R package for regression with multiple change points. *OSF Preprints*. (doi:10.31219/osf.io/fzqxv).
[33] Morris, Z.S., Vliet, K.A., Abzhanov, A. & Pierce, S.E. 2021 Developmental origins of the crocodylian skull table and platyrostral face. *Anat. Rec.* (doi:10.1002/ar.24802).
[34] Lenth, R.V. 2022 emmeans: estimated marginal means, aka least-squares mean. See https://cran.r-project.org/web/packages/emmeans.
[35] Collyer, M.L. & Adams, D.C. 2013 Phenotypic trajectory analysis: Comparison of shape change patterns in evolution and ecology. *Hystrix* **24**, 75-83. (doi:10.4404/hystrix-24.1-6298).
[36] Collyer, M.L. & Adams, D.C. 2021 RRPP: Linear model evaluation with randomized residuals in a permutation procedure. See https://cran.r-project.org/web/packages/RRPP.
[37] Revell, L.J. 2012 phytools: An R package for phylogenetic comparative biology (and other things). *Methods Ecol. Evol.* **3**, 217–223. (doi:10.1111/j.2041-210X.2011.00169.x).
